# Targeted Screening and Identification of Chlorhexidine as a Pro-myogenic Circadian Clock Activator

**DOI:** 10.1101/2023.01.18.524648

**Authors:** Tali Kiperman, Weini Li, Xuekai Xiong, Hongzhi Li, David Horne, Ke Ma

## Abstract

**Background:** Circadian clock is an evolutionarily-conserved mechanism that exerts pervasive temporal control in stem cell behavior. This time-keeping machinery is required for orchestrating myogenic progenitor properties in regenerative myogenesis that ameliorates muscular dystrophy. Here we report a screening platform to discover circadian clock modulators that promote myogenesis, with the identification of chlorhexidine (CHX) as a clock-activating molecule with pro-myogenic activities.

**Methods:** A high-throughput molecular docking pipeline was applied to identify candidate compounds with a structural fit for a hydrophobic pocket within the key circadian transcription factor protein, Circadian Locomotor Output Cycles Kaput (CLOCK). Secondary biochemical screen for clock-modulatory activities of these molecules were preformed, together with functional validations of myogenic regulations to identify modulators with pro-myogenic properties.

**Results:** CHX was identified as a clock activator that promotes distinct aspects of myogenesis. CHX activated circadian clock that reduced cycling period length and augmented amplitude. This action was mediated by the targeted CLOCK structure via augmented interaction with heterodimer partner Bmal1, leading to enhanced CLOCK/Bmal1-controlled transcription with up-regulation of core clock genes. Consistent with its clock-activating function, CHX displayed robust effects on stimulating myogenic differentiation in a clock-dependent manner. In addition, CHX augmented the proliferative and migratory activities of myoblasts.

**Conclusion:** Our findings demonstrate the feasibility of a screening platform to discover clock modulators with myogenic regulatory activities. Discovery of CHX as a pro-myogenic molecule could be applicable to promote regenerative capacities in ameliorating dystrophic or degenerative muscle diseases.

## Introduction

The circadian clock, driven by a transcriptional feedback loop coupled with translational and posttranslational regulations, generates the ~24-hour rhythm in physiology and behavior (1, 2). This evolutionarily-conserved time-keeping mechanism imparts pervasive temporal control in diverse biological processes, including metabolic regulation, cell cycle and stem cell behavior (3-8). Disruption of circadian clock regulation, increasingly prevalent in a modern lifestyle, leads to the development of metabolic disorders (9, 10), a myriad types of cancer (11) and dysregulation of tissue remodeling (6-8). Therapeutic targeting of circadian clock and its biological output pathways may have potential applications in metabolic disorders, cancer treatment and prevention, or dystrophic muscle diseases (6, 11-16). This spurred the current interests to identify clock-modulating molecules that may have potential utilities in these disease conditions (17-19), although therapeutic agents targeting the clock have yet to be discovered for diseases involving stem cells in tissue remodeling processes.

The molecular machinery that drives circadian rhythm, the molecular clock, is composed of a transcriptional/translational negative feed-back circuitry (2). The key transcription activators, CLOCK (Circadian Locomotor Output Cycles Kaput) and Brain and Muscle Arnt-like 1 (Bmal1), heterodimerize *via* the Per-Arnt-Sim (PAS) domain to form a transcription complex that drives clock gene transcription (20). Clock repressor proteins, Cryptochrome (Cry1 & 2) and Periods (Per1-3), are direct transcription targets of CLOCK/Bmal1, thereby constituting a negative transcriptional feed-back arm by inhibiting CLOCK/Bmal1 activity. CLOCK/Bmal1 also activates nuclear receptors, Rev-erbα and RORα, to exert positive and negative regulations via RORE response element to generate Bmal1 transcriptional oscillation, a mechanism that re-enforces the robustness of the clock (21, 22). Additional post-transcriptional and post-translational regulatory controls are involved to complete the molecular clock circuit that drives the 24-hour oscillations in gene expression, physiology and behavior (1). Recently, various components of the molecular clock circuit have been targeted for pharmacological modulations, although small molecule clock modulators discovered so far target the negative regulators such as Rev-erbα and Cry (12, 13, 23-25). To date, molecules directly modulating the key driver of the molecular clock, CLOCK/Bmal1-mediated transcriptional activation, remains to be identified.

Accumulating studies indicate that circadian clock exert important temporal control in distinct aspects governing stem cell behaviors (7, 8, 26, 27). In skeletal muscle, in addition to maintenance of nutrient metabolism and structural integrity, the circadian clock is intimately involved in modulation of muscle stem cell myogenic properties in tissue growth processes (6, 28, 29). A functional muscle clock is required to facilitate metabolic fuel switch from fat oxidation to glucose utilization as a nutrient sensor that determines insulin sensitivity by responding to feeding signals (30), and this metabolic role of the muscle clock was also highlighted by recent findings of augmented metabolic rhythm by exercise regimen (29, 31). Our previous studies revealed that the clock is required to orchestrate myogenic progression of muscle stem cells (32-35). The concerted clock regulation to facilitate myogenic progenitor differentiation into mature multi-nucleated myotubes involves both the positive arm and negative regulatory arms of the circadian clock transcriptional feed-back regulatory circuit. Bmal1 deficiency impairs myogenic differentiation and regenerative myogenesis, whereas ablation of its transcription repressor Rev-erbα promotes these processes (32, 33, 36). In addition, Per1/Per2 and Cry2, key regulators within the negative molecular clock loop, modulates myogenesis that impacts muscle regeneration (34, 35). Moreover, loss of *Bmal1* resulted in obesity with reduced muscle mass (32, 37). Thus, targeting the circadian clock function, based on its modulation of myogenic progenitors in skeletal muscle, may have potential therapeutic utilities to promote regenerative capacity in dystrophic or degenerative muscle diseases.

Despite recent development, current efforts in discovering clock modulators are focused on metabolic diseases and cancer therapy. Agonists for ligand-binding nuclear receptors Rev-erbα and ROR displayed metabolic benefits (24, 38-40), while Cry-stabilizing molecules may have potential utility in neuroblastoma (13, 14, 25). To date, small molecules directly targeting the key transcriptional driver of the core clock loop, the CLOCK protein, has yet to be uncovered. In the current study, we conducted a screen for compounds specifically targeting the CLOCK protein to modulate circadian clock activity and myogenesis. Applying this screening platform, we identified chlorhexidine (CHX) as a novel clock activator with pro-myogenic activities that may have potential muscle disease utilities.

## Materials & Methods

### Cell culture

The sources of cell lines used in the study were listed in Suppl. Table 1. Cells were maintained at 37°C in 10% Fetal bovine serum (FBS) (Cytiva), 1% Penicillin-Streptomycin-Glutamine (PSG) (Gibco – Thermo Fisher) Dulbecco’s Modified Eagle Medium (DMEM) (Gibco – Thermo Fisher) and removed from culture plates for experimentation using 0.25% Trypsin –EDTA (Gibco – Thermo Fisher CN: 25200072). 2% FBS supplemented DMEM was used for differentiation of 80–90% confluent cultures on C2C12, C2C12 Bmal1 KD and C2C12 SC cell lines.

### All-around molecular docking-based screening pipeline

Briefly, the protein crystal structure of CLOCK was obtained from RCSB Protein Data Bank (PDB 4f31 (20). An in-house developed LiVS (Ligand Virtual Screening Pipeline), as previously described (41, 42) was employed to screen the NCI Developmental Therapeutics Program (DTP) compound library (containing about 260,000 compounds) and available FDA library in-silico to identify structural hits. LiVS method is a multiple-stage and full-coverage pipeline for ligand screening that utilizes the three precision modes (i.e., HTVS, high-throughput virtual screening; SP, standard precision; and XP, extra precision) of Schrodinger Glide software (43) for docking analysis. First, a HTVS precision mode, which is fast but less accurate, was implemented to dock the entire library. The 10,000 top-ranked compounds were next docked and scored by the SP mode. Then the 1,000 top-ranked compounds from SP precision docking were re-docked and re-scored by the XP mode. The 1,000 compounds were further analyzed and filtered by Lipinski’s rule of five (44), HTS frequent hitter (PAINS) (45), protein reactive chemicals such as oxidizer or alkylator (ALARM) (46), and maximized the molecule diversity by using UDScore (Universal Diversity Score, developed by us to measure library diversity which is independent of library size). Based on the virtual screening pipeline, we requested the top 266 compounds from NCI DTP and obtained 83 available for secondary functional screening.

### Luminometry, real-time bioluminescence recording and data analysis

U2OS cells containing a stable Period2 promoter-driven luciferase (*Per2::dLuc*) reporter (47, 48) were seeded at 4×10^5^ concentration on 24 well plates and treated at 90% confluence with 1ml fresh explant medium with compounds at indicated concentration and DMSO control. The optimal seeding density was chosen based on 90% confluence at 24 hours after seeding. Fresh explant medium contains 50% 2xDMEM buffer stock [DMEM powder (Thermo Fisher CN: 12800-017), Sterile MilliQ water, 1M pH7 HEPES (Thermo Fisher CN:15630080) and 7.5% Sodium Bicarbonate (Thermo Fisher CN: 25080094)], 10% FBS (Cytiva), 37.9% Sterile MiliQ water, 1% PSG 100X (Gibco – Thermo Fisher CN: 10378016), 0.1% Sodium Hydroxixe (NaOH) 100mM (Fisher Chemical CN: S318-500), 1% XenoLight D-Luciferin Monopotassium Salt Bioluminiscence Substrate 100mM (PerkinElmer CN: 122799). Plates were sealed with plastic film and placed inside LumiCycle96 (ActiMetrics) in a bacterial incubator at 36^0^C, 0% CO2, and maintained for 7 days of Per2-luciferase luminescence recording. Measurement of bioluminescence rhythms from *Per2::dLuc* U2OS reporter line was conducted by LumiCycle 96 luminometer, as described (47, 49, 50). Briefly, luminescence from each well was measured for

∼70 s at intervals of 10 min and recorded as counts/second. Raw and subtracted results of real-time bioluminescence recording data for 6 days were exported, and data was calculated as luminescence counts per second. LumiCycle Analysis Program (Actimetrics) was used to determine clock oscillation period, length amplitude and phase. Briefly, raw data following the first cycle from day 2 to day 5 were fitted to a linear baseline, and the baseline-subtracted data (polynomial number = 1) were fitted to a sine wave, from which period length and goodness of fit and damping constant were determined. For samples that showed persistent rhythms, goodness-of-fit of >80% was usually achieved.

### Plasmids and shRNA

pcDNA3-BMAL1-His (AddGene CN: 31367) were purchased from the Addgene. PcDNA3.0-6XMyc-Clock and pcDNA3.0-3XFLAG-cry2 were generated by PCR amplification and sub-cloned in pCDNA3.0–6XMyc or pCDNA3.0–3XFLAG vector separately. The primers for PCR amplifications are: CLOCK-s: GCGGCCGCATGGTGTTTACCGTAAGC, CLOCK-as: CTCGAGTTCAGCCCTAACTTCTGCA; Cry2-s: GCGGCCGCATGGCGGCGGCTGCTGTGGT Cry2-as: CTCGAGTCAGGAGTCCTTGCTTGCT. The cysteine267 mutation of *CLOCK* gene was generated using site-directed mutagenesis. The primer sequences to generate the mutations are: *CLOCK*-C267A-s: TTCATCAAGGAAATGGCTACTGTTGAA, *CLOCK*-C267A-as: GCCATTTCCTTGATGAACTGAGGTG. The short hairpin RNAs of CLOCK used to generate CLOCK stable KD U2OS cells was purchased from Sigma-Aldrich (TRCN0000018976) and cloned into PLKO.1 puro vector.

### Generation of stable cell lines

Stable clones of U2OS cell line containing control or sh*CLOCK* were generated by lentiviral transduction and stable selection. Briefly, HEK293A were transfected with lentiviral packaging plasmids (psPAX2 and pMD2.G) and lentivirus vectors PLKO.1 or PLKO.1-sh*CLOCK* using PEI Max. At 48 hours post-transfection, lentiviruses were collected through 0.45um filter to remove the cell debris. U2OS cell line was infected by the lentivirus medium supplemented with polybrene (Santa Cruz, CN: SC-134220). 24 hours following lentiviral infection, stable cell lines were selected in the presence of 1 μg/ml puromycin.

### Transient transfection luciferase assay

Luciferase assays were performed as previously described (51). Briefly, cells were seeded at 4×10^5^ concentration on 24 well plates. At 90% confluence cell transfection was performed using Opti-MEM™ Reduced Serum Medium (Thermo Fisher CN:31985062) with PGL2, PRL, PcDNA3.0-6XMyc-CLOCK (AddGene CN: 47334) pcDNA3.0-BMAL1-His (AddGene CN: 31367), pcDNA3.0-3XFLAG-Cry2 plasmids and PEI max 40k (Polysciences Inc CN: 24765-1). 24 hours after transfection, cells were treated with desired compounds and controls in 10% FBS 1% PSG DMEM media for desired times. Cells were extracted with 1XPLB solution provided in the Dual-Luciferase® Reporter Assay Kit (Promega CN: E1910) and transferred to 96-well black plate to be treated with LARII and Stop & Glo components according to protocol of the Dual-Luciferase® Reporter Assay Kit (Promega CN: E1910). Luminescence was measured using a microplate reader (TECAN infinite M200pro). The mean and standard deviation values were calculated for each well and graphed.

### RNA extraction and quantitative reverse-transcriptase PCR analysis

Cells extracted with TRIzol™ Reagent (Ambion CN: 15596018) were collected. RNA extraction was performed with Chloroform (Fisher Chemical CN: C298-500) and Isopropanol (Fisher Chemical CN: A451SK-4) extraction followed by two purification steps with 75% Ethanol (Fisherbrand CN: HC-800-1GAL) and elution in sterile MilliQ water. RNA concentration was measured using Nanodrop spectrophotometer (Thermo Fisher). Following RNA extraction, reverse transcription was carried out using RevertAid RT Reverse Transcription Kit (Thermo Fisher CN: K1691) and run on SimpliAmp Thermal Cycler (Thermo Fisher CN: A24811) according to the kit procedure. Quantitative PCR was performed using PowerUp™ SYBR™ Green Master Mix (Thermo Fisher CN: A25742) using a dilution of 5:2:1 master mix, RNA and H2Omq according to the protocol on ViiA 7 Real-Time PCR System (Applied Biosystems). Primer sequences used are specified in Suppl. Table 2. Relative expression levels were determined using the comparative Ct method to normalize target genes to 36B4 internal controls.

### Immunoblot analysis

Total protein (20-30 ug) was extracted using standard immunoprecipitation lysis buffer (3% NaCl, 5% Tris-HCl, 10% Glycerol, 0.5% Triton X-10 in Sterile MilliQ water) and resolved on 10% SDS-PAGE gels followed by western blotting on Immun-Blot PVDF membranes (Bio-rad). Antibodies used were diluted in 5% milk (Labscientific CN: M0841). Primary and secondary antibodies with dilutions used are described in Suppl. Table 3. Membranes were washed with TBS-T (10%TBS, 0.1% Tween 20, Fisher BioReagents CN: BP377-500). Images were developed using SuperSignal West Pico PLUS Stable Peroxide Solution (Thermo Scientific CN: 1863095) and SuperSignal West Pico PLUS Luminol Enhancer Solution (Thermo Scientific CN: 1863094) on chemiluminescence imager (GE Healthcare Bio-Sciences AB Amersham Imager 680).

### Immunoprecipitation

HEK293A cells were transfected with pcDNA3.0-BMAL1-His, PcDNA3.0-6XMyc-CLOCK or PcDNA3.0-6XMyc-CLOCKC267A using Polyethylenimine MaX based on the manufacturer’s instructions. 24 hours after transient transfection, cells were treated with indicated compounds for 8 hours and homogenized in lysis buffer (50 mM Tris pH7.5, 150mM NaCl 0.5% Triton X-100, 5% glycerol and protease inhibitor). Immunoprecipitation was performed overnight using Anti-c-Myc Agarose beads (Thermo Fisher CN: 20168), following by washing for three times with lysis buffer. Then the samples resolved by SDS-PAGE and immunoblotted with indicated antibodies as described in Suppl. Table 3.

### Myogenic differentiation of C2C12 myoblasts

C2C12, C2C12 Bmal1 KD and C2C12 Scrambled control (SC) myoblasts were seeded at 1-2×10^5^ density on 6 wells plates and maintained in 10% FBS until 90% confluency. 2% FBS DMEM media was used to induce differentiation as day 0, together with indicated compounds or DMSO control. Phase-contrast images were taken daily to monitor morphological progression, and proteins and RNA samples were collected on days indicated.

### Myosin heavy chain immunofluorescence staining

C2C12, C2C12 Bmal1 KD and C2C12 Scrambled control (SC) myoblasts were seeded at Day -1 at 2×10^5^ concentration on 12 wells plates and maintained in 10% FBS until 80% confluency and treated with indicated compounds and controls in 2% FBS DMEM media to induce differentiation on Day 0. On days 3, 5 and 7 cells were fixed with 4% paraformaldehyde (from 37% Formaldehyde Baker Analyzed) and kept at 4°C in PBS. Cells were permeabilized with 0.5 % Triton X-100 (Fisher Bioreagents CN: 9002-93-1) in PBS and blocked with 1% BSA (Fisher Bioreagents CN:9048-46-8) in PBS. Primary antibodies (Suppl. Table S4) were diluted in washing solution of 0.01% Triton X-100 (Fisher Bioreagents CN: 9002-93-1) in PBS and left overnight at 4°C. Cells were then incubated with secondary antibodies (Table 4) in washing solution at room temperature for 1h. 1 ug/ml 4’,6’-diamidine-2’-phenylindole dihydrochloride (DAPI) was used to label nuclei. Images were acquired using Echo Revolve fluorescence microscope at 10X magnification. To quantify the proliferation of myoblast, when needed, cells were treated with 1 μg/ml of EdU for 3 hours before fixation.

### EdU proliferation assay

Cells were seeded at 0.2×10^5^ concentration and treated with desired compounds and controls overnight. The seeding density was decided after trying several densities and choosing the one that after 24hs lead to visible, not clumping cells. 10 μM 5-ethynyl-2′-deoxyuridine (EdU) incorporation was performed for 4 hours according to the protocol. Detection of EdU was performed with Click-iT EdU Imaging Kit with Alexa Fluor 488 (Invitrogen). 1 ug/ml 4’,6’-diamidine-2’-phenylindole dihydrochloride (DAPI) was used to label for nuclei. Images were acquired using Echo Revolve fluorescence microscope at 20X magnification. Total number of EdU+ cells was counted from 10 representative fields at 20X and the rate of proliferation was calculated as percentage of EdU^+/^DAPI.

### Wound healing assay

Cells were seeded at 12×10^5^ concentration on 6 well plates until 90% coverage and treated with compounds as indicated and controls while maintained 10% FBS 1% PSG DMEM. Scratch wounds were made with disposable cells scrapers and pictures were taken at hours 0, 4, 8, 24 and 48 following scratch wound. The results were measured as percentage coverage over original wound area based on image quantification. Area of wound closure was measured using Image J software.

### Statistical analysis

Experiments results were analyzed and graphed using PRISM, and data were presented as mean+ SD. Each experiment was repeated at minimum twice to validate the result. Sample sizes were indicated for each experiment in figure legends. Two-tailed Student’s t-test or One-way ANOVA with post-hoc analysis for multiple comparisons were performed as appropriate as indicated. P<0.05 was considered statistically significant.

## Results

### Screening pipeline for small molecule modulators of circadian clock

Recent therapeutic targeting of circadian clock mostly focused on repressor proteins in the molecular clock feed-back loop, including nuclear Rev-erbs and Cryptochromes. To identify small molecules targeting the core clock transcription activator, CLOCK, we conducted an in-silico molecular docking-based screening of chemical libraries from NCI Developmental Therapeutic Program (DTP) with (∼275,000 compounds) and the FDA library (4,086 compounds). Crystal structure of CLOCK protein, PDB 4f31 (20), was used for molecular docking modeling for high-throughput virtual screening strategy to identify compounds with structural fit (42), with the screening strategy pipeline shown in **Fig. 1A**. All-around docking (ADD) analysis identified hit molecules with ranked structural fit for a deep hydrophobic pocket we discovered, within the well-defined PAS-A domain of CLOCK protein that mediates heterodimerization with its obligatory partner Bmal1 (20). Compounds were ranked based on Glide Score calculation that predicts potential strength of binding with predicted hydrophobic, polar, or hydrogen bond interactions with key residues within this structure. 266 compounds were identified as hits in these libraries based on Glide Score ranking of ≤3.5, and the docking poses were shown in **Fig. 1B**. Out of these initial hits *via* virtual screening, we were able to obtain 84 molecules for secondary screening through experimental validation for clock-modulatory activity. Employing a gold-standard clock activity assay *via* a U2OS luciferase reporter cell line containing a Period 2 gene promoter-driven luciferase construct (*Per2::dLuc*) (47), this secondary screening identified 8 molecules with significant clock modulatory activity. Chlorhexidine (CHX) was the only molecule that exhibited clock-activating properties. CHX displayed a strong GS score of ∼19.5, with a tight docking pose within the targeted CLOCK hydrophobic pocket (**Fig. 1C and Fig. S1A**). Predicted interactions of CHX with the key amino acid residues lining the CLOCK pocket structure within 10-20A^0^ include charged, hydrophic, polar or hydrogen bonds were indicated (**Fig. 1D**), with the detailed analysis of theses interactions demonstrated in **Fig. S1B**.

**Figure 1.**
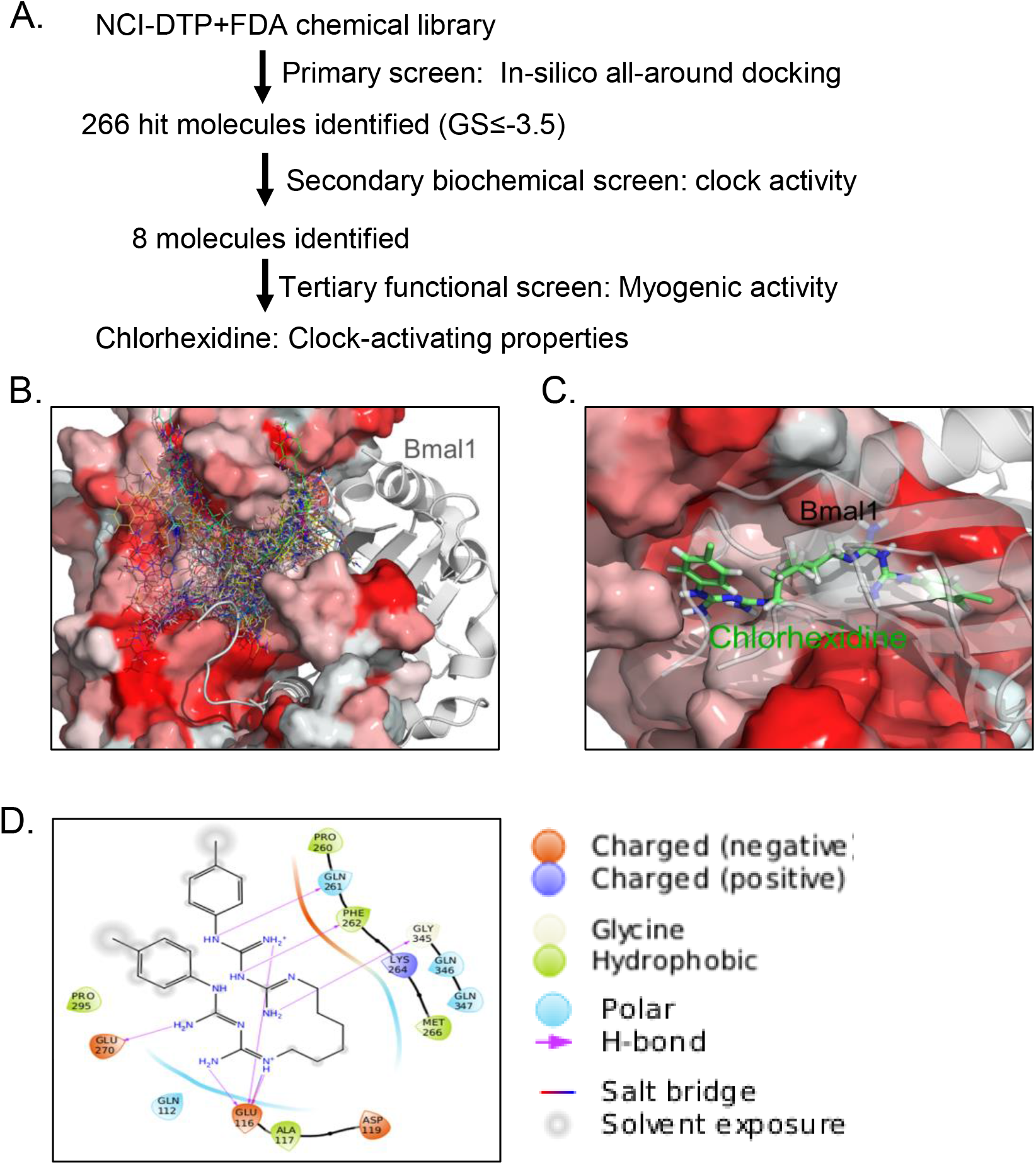
In-silico screening and identification of chlorhexidine as a circadian clock modulator. (A) Docking poses of 266 hit compounds identified via LiVS modeling targeting the CLOCK protein hydrophobic pocket that lies within CLOCK-Bmal1 interaction interface. Crystal structure of heterodimeric CLOCK (red) with Bmal1 (gray) is based on PDB: 4f3l. (B) Screening and biochemical validation pipeline for identification of clock modulators using NCI/DTP-FDA chemical libraries. (C) Docking pose of Chlorhexidine within the CLOCK protein hydrophobic pocket structure. CLOCK protein is shown in red and Bmal1 as gray. (D) Predicted interactions of Chlorhexidine with key amino acid residues of the CLOCK hydrophobic pocket.

### Identification of Chlorhexidine as a circadian clock activator

The activities of CHX on modulating key properties of circadian clock function, including oscillating period length and amplitude was determined *via* continuous monitoring of bioluminescence activity of the *Per2::dLuc* U2OS reporter cell line (47, 48). Cells were treated with CHX at indicated concentrations from 0.2 to 2 μM at start of the bioluminescence recording, as shown in original and baseline-subtracted plots for 6 days (**Fig. 2A & Fig. S2A**). Quantitative analysis revealed reductions of period length by CHX as compared to the DMSO control with the strongest effect observed at 1μM (**Fig. 2B**). This effect of CHX on period length shortening revealed its activity to activate clock leading to shorter cycles. In addition, CHX at 0.2 μM induced increased clock amplitude, but not at higher concentrations examined (**Fig. 2C**). Additional higher concentrations of CHX tested, at 5 and 10 μM, revealed toxicity with reduced amplitude (data not shown). To determine the direct effect of CHX on clock activation, we tested acute CHX treatment on CLOCK/Bmal1-mediated transcription via transient transfection to activate a Per2-luciferase reporter (48, 52). Bmal1 and CLOCK co-transfection induced Per2-luciferase activity by ∼4-fold, as expected (**Fig. 2D**). CHX treatment for 6 hours at 0.2 μM was sufficient to stimulated ∼26% higher levels of CLOCK/Bmal1-activated transcription with similar effects at higher concentrations. Addition of Cry2 was sufficient to suppress CLOCK/Bmal1-medaited transcriptional activation, as expected, while CHX showed moderate effect in increasing luciferase activity under Cry2 repression at 0.2 μM.

**Figure 2.**
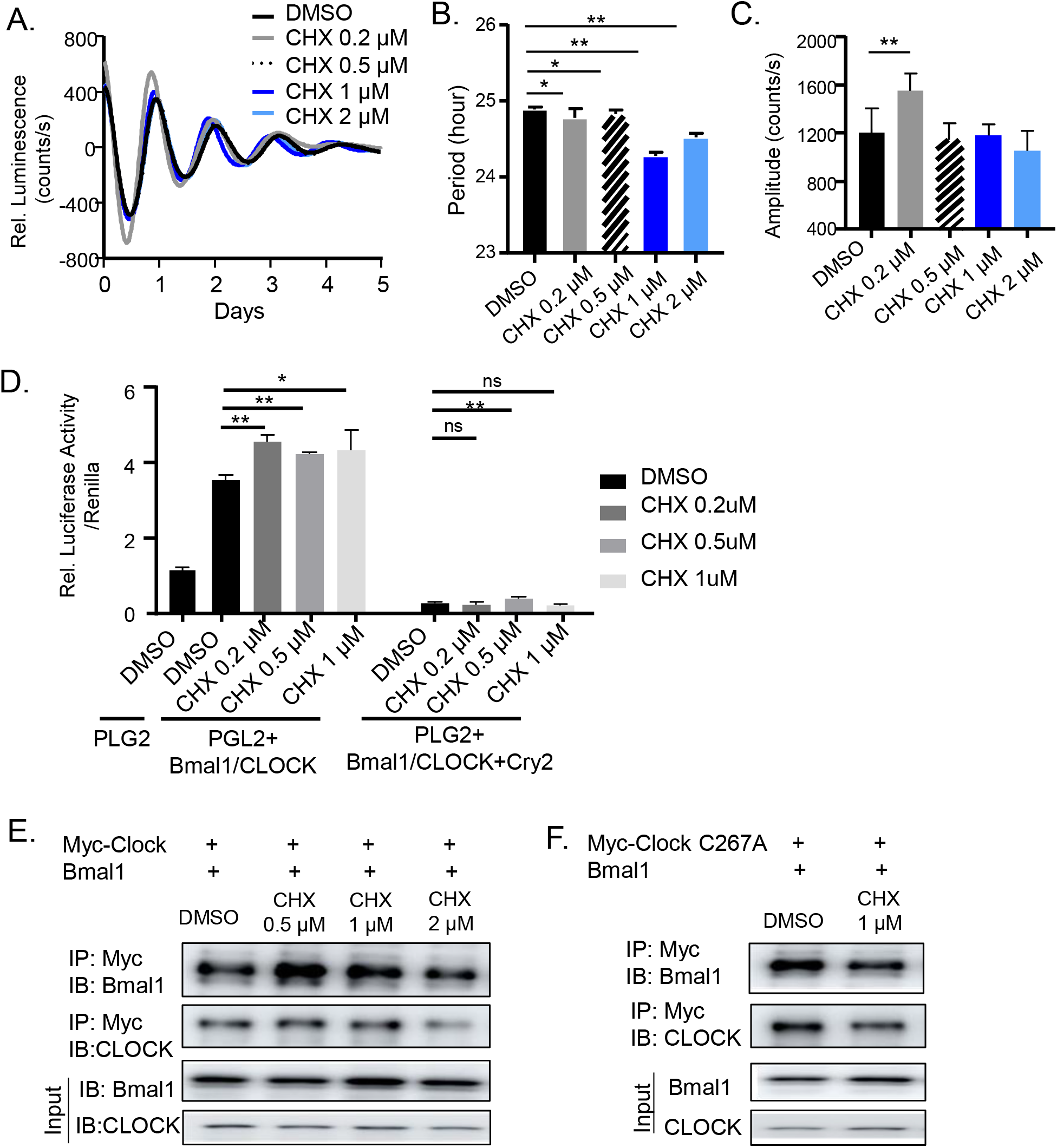
Biochemical characterization of chlorhexidine as a circadian clock activator. (A) Average tracing of bioluminescence activity monitoring of *Per2::dLuc* U2OS reporter cell line for 6 days using Lumicycle, with chlorhexidine treatment (CHX) at indicated concentrations as shown in baseline-adjusted plots. (B, C) Quantitative analysis of clock period length (B) and cycling amplitude (C) at CHX concentrations as indicated. Data are presented as Mean ± SD of n=4 replicates for each concentration tested, with four independent repeat experiments. *, **: p<0.05 and 0.01 CHX vs. DMSO by Student’s t test. (D) Transient transfection luciferase assay using a Per2-driven luciferase reporter in U2OS cells with indicated CHX concentration. PGL2: empty vector control. N=5 for each concentration tested. *, **: p<0.05 and 0.01 CHX vs. DMSO by Student’s t test. (E, F) Co-immunoprecipitation analysis of Bmal1 interaction with Myc-tagged CLOCK protein (E), or Myc-tagged C267A mutant CLOCK protein (F), with or without CHX treatment for 16 hours at indicated concentrations in 293T cells. Uncropped gels are shown in Fig. S5.

Given our screening strategy by targeting the CLOCK protein hydrophobic pocket involved in heterodimerization with Bmal1, compounds with a structural fit may modulate this interaction to influence CLOCK/Bmal1 transcriptional activity. We thus tested whether CHX activation of clock is mediated through promoting CLOCK/Bmal1 interaction to enhance transcription. Via co-immunoprecipitation to detect CLOCK binding with Bmal1, CHX at 0.5 and 1 μM were sufficient to augment this interaction with increased Bmal1 level immunoprecipitated by Myc-CLOCK, although 2 μM does not further enhance this activity (**Fig. 2E**). To test whether this effect of CHX on CLOCK/Bmal1 interaction depends on the targeted CLOCK pocket structure, we mutated a key residue, Cysteine 276, to Alanine. While this mutation did not affect CLOCK/Bmal1 heterodimer association as compared to wild-type CLOCK with efficient Bmal1 detection in immunoprecipitated protein complex, CHX failed to increase the interaction between Bmal1 and the CLOCK C267 mutant, suggesting that CHX effect is dependent on the targeted CLOCK structure **(Fig. 2F**). Collectively, these results reveal that CHX is a novel circadian clock activator and functions by promoting CLOCK/Bmal1 interaction to activate transcription.

### Chlorhexidine induces clock gene expression

Based on findings of CHX in stimulating CLOCK/Bmal1-mediated transcription, we determined its potential effect on inducing clock gene expression in U2OS cells. CHX at 0.1 and 0.2 μM were sufficient to induce CLOCK protein levels, with elevated Bmal1 expression at 0.2 and 0.5 µM (**Fig. 3A** & **3B**). Rev-erbα and Period 2 (Per 2), direct transcription targets of CLOCK/Bmal1 in the core clock circuit, were also induced by CHX in a largely dose-dependent manner. Moderately attenuated effect of CHX was observed at 0.5 µM for these proteins except Rev-erbα. Consistent with these findings, further analysis of *CLOCK* and *Bmal1* transcripts revealed marked up-regulation by CHX (**Fig. 3C**). Additional components of molecular clock that are under direct transcriptional control of CLOCK/Bmal1, including *Dbp, Nr1d1, Nr1d2* and *Cry2*, were also induced by CHX treatment (**Fig. 3D** & **3E**), suggesting increased CLOCK/Bmal1 transcriptional activity.

**Figure 3.**
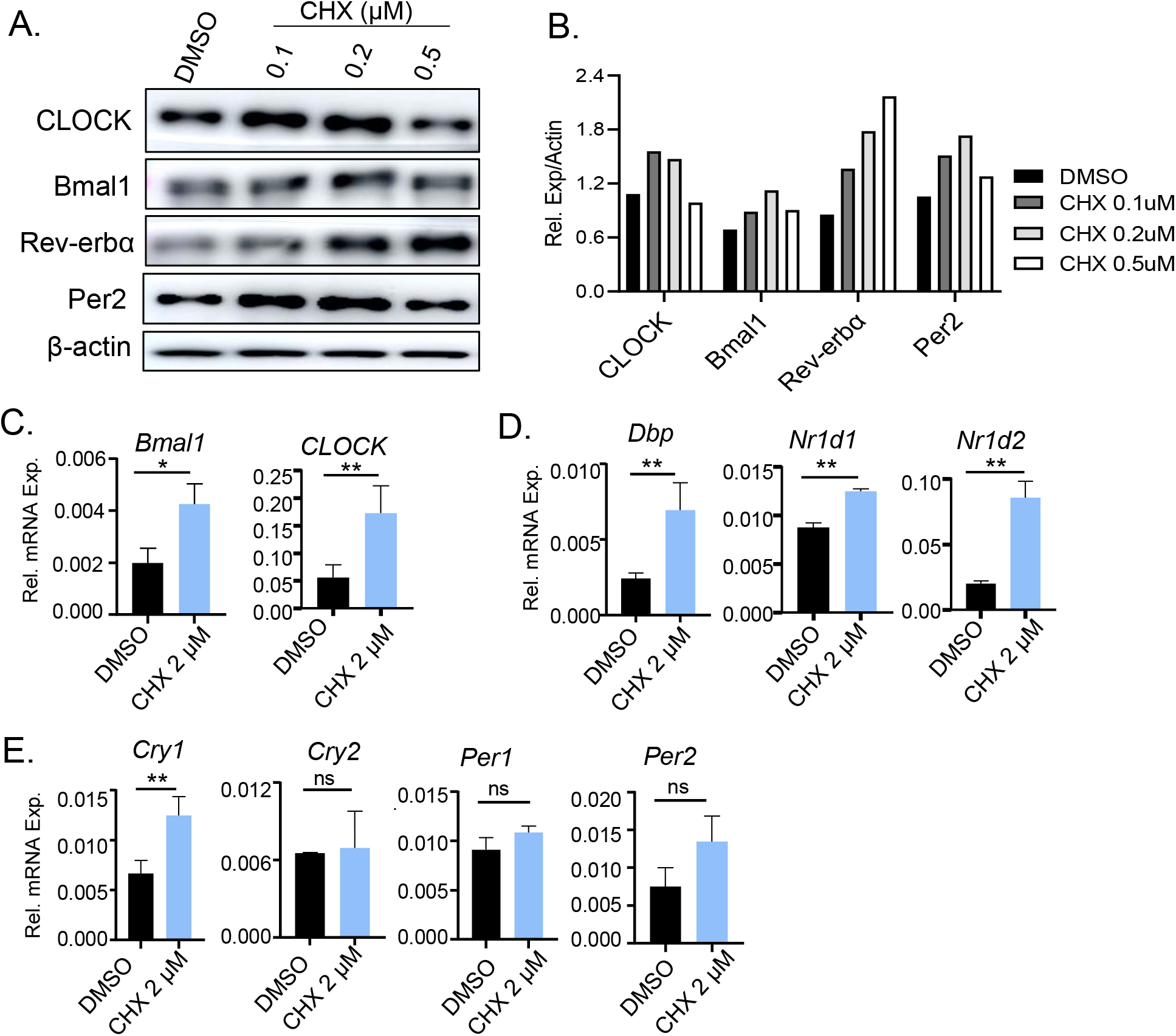
Effect of chlorhexidine on core clock gene expression in osteosarcoma cells. (A, B) Immunoblot analysis of core clock protein expression with CHX treatment at indicated concentrations for 24 hours in U2OS cells (A) with quantitative analysis (B). Each lane represents pooled samples of 3 replicates. Uncropped gels are shown in Fig. S6. (C-E) RT-qPCR analysis of clock genes, including core clock activators (C), Bmal1/CLOCK target genes (D) and clock repressor genes (E), after CHX treatment for 24 hours in U2OS cells. Data are presented as Mean ± SD of n=3 replicates with two independent repeat experiments. *,**: p<0.05 and 0.01, CHX vs. DMSO by Student’s t test.

Circadian clock components coordinate the progression of myogenic differentiation (6, 32, 33). Using C2C12 mouse myoblasts as a cellular model, we determined whether CHX modulates clock activity in myogenic progenitors. Through myoblast differentiation for day 2, 4 and 6 days, CHX at 0.5 µM robustly stimulated CLOCK and Bmal1 protein levels, together with their transcriptional targets Rev-erbα and Cry2 (**Fig. 4A** & **Fig. S2B**). Bmal1, Rev-erbα and Cry2 protein also responded to 0.2 µM CHX at day 6. The augmented clock protein expression by CHX was further corroborated by induction of these transcripts in a largely dose-dependent manner (**Fig. 4B**). Furthermore, we tested whether CHX induction of molecular clock gene expression is dependent on a functional clock, using myoblasts containing stable shRNA silencing of *Bmal1* (BMKD) as compared to cells with scrambled control (SC)(32). As expected, CHX induced *Clock, Bmal1* and their transcription targets, *Nr1d1* and *Per2* in SC myoblasts (**Fig. 4C**) at day 4 of myogenic differentiation, while this effect was completely abolished in BMKD cells with loss of Bmal1 function. Thus, indeed CHX effect on activating clock gene transcription is dependent on a functional clock.

**Figure 4.**
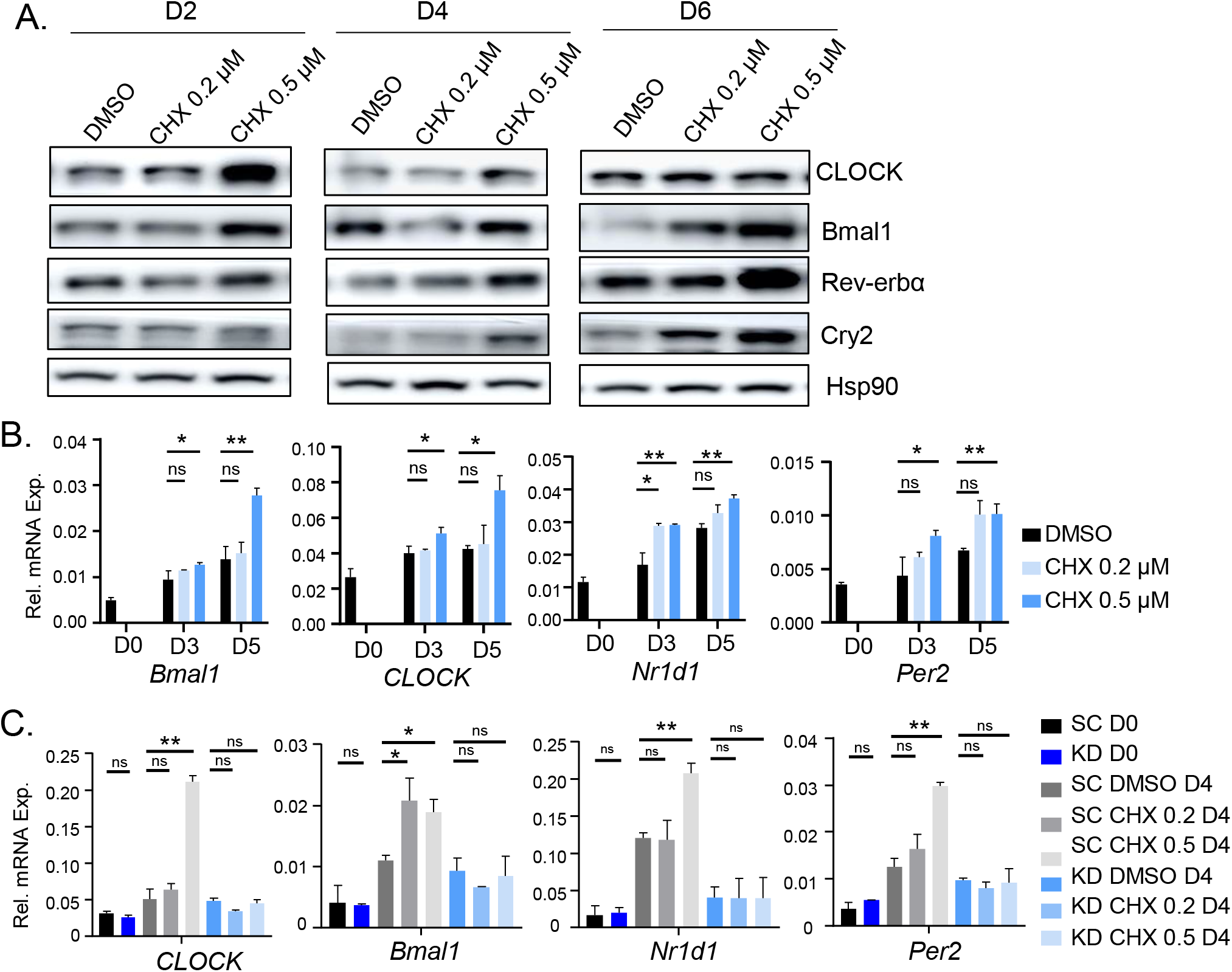
Effect of chlorhexidine on clock gene regulation in C2C12 myoblasts. (A) Immunoblot analysis of CHX effect on clock protein expression in C2C12 myoblasts during day 2, 4, and 6 of myogenic differentiation at indicated concentration. Each lane represents pooled samples of 3 replicates. Uncropped gels are shown in Fig. S7. (B) RT-qPCR analysis of CHX treatment on expression of clock genes at day 0, 3 and 5 of myogenic differentiation. (C) RT-qPCR analysis of expression of clock genes in C2C12 myoblasts with stable expression of scrambled control shRNA (SC) or Bmal1 shRNA (BMKD) in response to CHX treatment before and at day 4 of C2C12 myogenic differentiation. Data are presented as Mean ± SD of n=3 replicates with three independent repeat experiments. *, **: p<0.05, and 0.01 CHX vs. DMSO by Student’s t test.

### Clock-dependent effect of Chlorhexidine on promoting myogenesis

Our previous studies demonstrated that the essential clock activator, Bmal1, promotes myogenesis *via* direct transcriptional control of components of the Wnt signaling cascade(15, 32, 36). We postulated that CHX, as a clock-activating molecule, may display pro-myogenic activity in myoblast differentiation. As shown by phase-contrast images, normal C2C12 myoblasts differentiate efficiently into mature multi-nucleated myotubes upon 2% serum induction for 9 days (**Fig. 5A**). In comparison, CHX treatment at 0.2 or 0.5 µM markedly accelerated the morphological progression of myotube formation, with abundant mature myotubes formed at early stage of differentiation at day 3 when barely visible mature myotubes were detected in DMSO-treated cells. Myotube formation in day 3 CHX-treated cells were nearly complete, and comparable to the differentiation observed at day 6 controls, indicating enhanced myocyte maturation by CHX. This effect was maintained throughout the differentiation time course, with CHX-treated cells displaying more advanced myotube elongation at day 6 and 9 as compared to controls. However, higher concentrations of CHX at 2 and 5 μM appeared to be toxic to myoblasts with significant cell death during differentiation. This pro-myogenic activity of CHX was further demonstrated by myosin heavy chain (MyHC) immunofluorescence staining to identify myocytes during differentiation. Consistent with the advanced morphological progression, CHX treatment increased numbers of MyHC-positive mature myotubes at day 5 and day 7 of differentiation, and 0.5 µM CHX induced notable myotube hypertrophy at Day 7 (**Fig. 5B**). This effect was evident as early as at day 3 of differentiation (**Fig. S3**). In line with the augmented myogenic differentiation, CHX led to early induction of myogenic factors, Myf5 and Myogenin proteins, at day 3 and day 5 along the differentiation time course (**Fig. 5C**). *Myf5, Myod1* and *embryonic myosin heavy chain (eMyhc)* transcripts also displayed significant up-regulations by 0.5 µM CHX with a tendency toward higher expression at 0.2 µM (**Fig. 5D**). Mature myocyte markers, Myogenin and myosin light chain expression were similarly induced by CHX (**Fig. S3C**).

**Figure 5.**
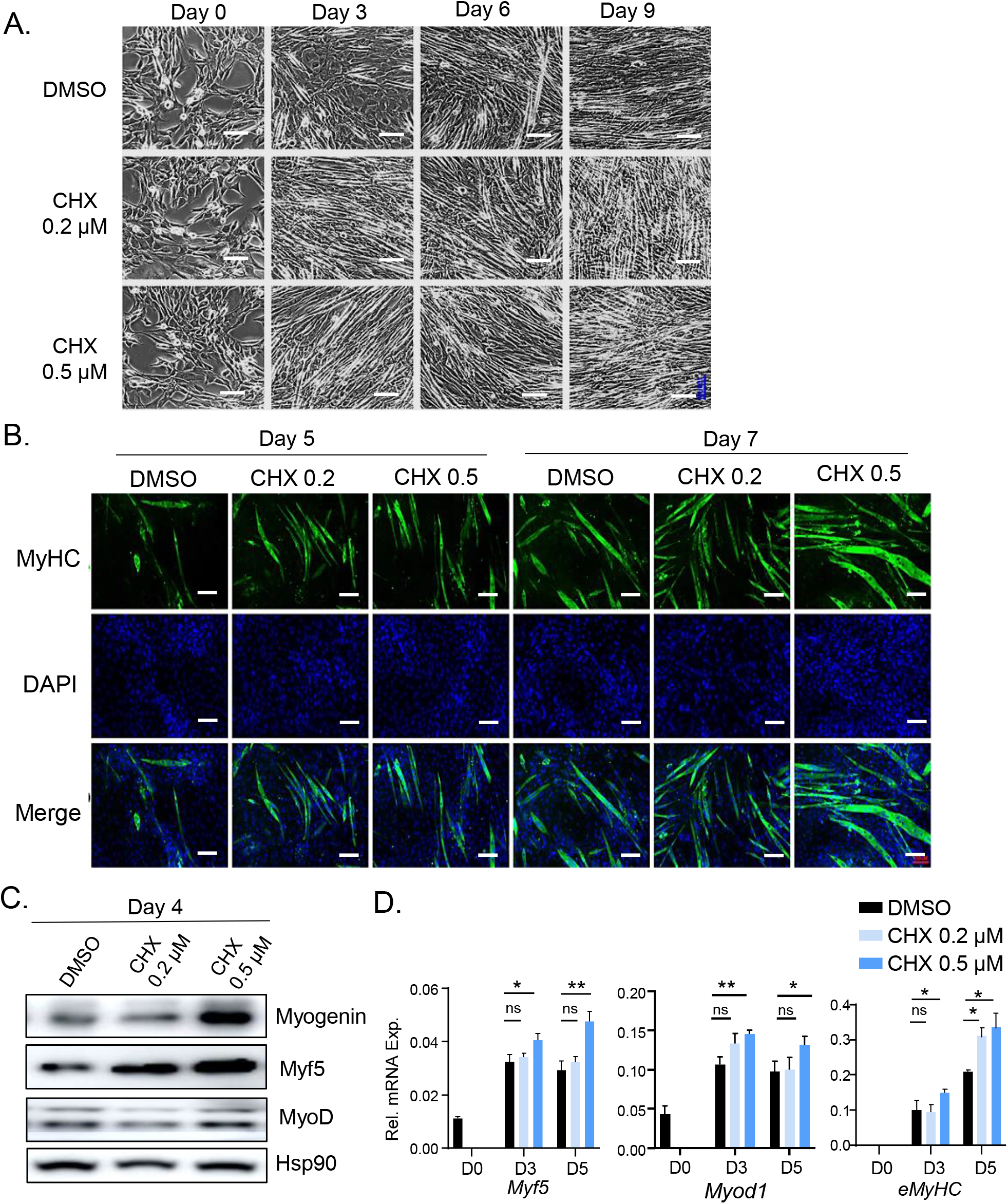
Effect of chlorhexidine on promoting C2C12 myogenic differentiation. (A) Representative phase-contrast images of C2C12 morphology on day 0, 3, 6 and 9 of myogenic differentiation with CHX treatment. (B) Representative images of myosin heavy chain (MyHC) immunofluorescence staining at day 5 and day 7 of myogenic differentiation. Scale bar: 100 µm. (C, D) Immunoblot analysis (C), and RT-qPCR analysis of myogenic gene expression (D) at indicated CHX concentrations during C2C12 myogenic differentiation. Uncropped gels are shown in Fig. S8. Data are presented as Mean ± SD of n=3 replicates with three independent repeat experiments. *,**: p<0.05 or 0.01 CHX vs. DMSO by Student’s t test.

Next, we examined the pro-myogenic activity of CHX using BMKD myoblasts to determine whether this effect is dependent on clock modulation. Due to inhibition of Bmal1, myogenic differentiation of BMKD myoblasts were impaired as compared to SC control (**Fig. 6A**), as previously reported(32). CHX stimulated myotube formation of SC myoblasts similarly as observed for the parental C2C12 myoblasts, albeit the rate of myogenic progression was attenuated (**Fig. 6A**). In contrast, CHX failed to augment BMKD myoblast differentiation, indicating that its pro-myogenic activity is clock-dependent. In the KD cells, MyHC staining for mature myocytes revealed a similar finding of loss of CHX effect on enhancing maturation of myocytes at day 5 of differentiation as compared to its robust effect in SC myoblasts (**Fig. 6B**). Gene expression analysis further confirmed the clock-dependent effect of CHX on inducing the myogenic program, with loss of CHX induction of *Myf5* and *eMyhc* in Bmal1-deficient myoblasts (**Fig. 6C**). Clock exert direct transcriptional control on genes involved in disparate steps of Wnt signaling pathway during myogenesis(32). We thus examined the known transcriptional targets of clock in this pathway and found that CHX was able to stimulate upstream Wnt ligands including Wnt10a and Wnt10b, as well as the Wnt receptor Frizzled 5 (Fzd5). In addition, CHX induced the expression levels of the key effector of Wnt signaling β-catenin, transcription factor TCF3 and the Wnt target gene Axin 2 in SC myoblasts, while this effect was abolished in Bmal1-deficient cells (**Fig. 6D** & **6E**), indicating clock-dependent actions of CHX in activating a key signaling mechanism that drives myogenic differentiation.

**Figure 6.**
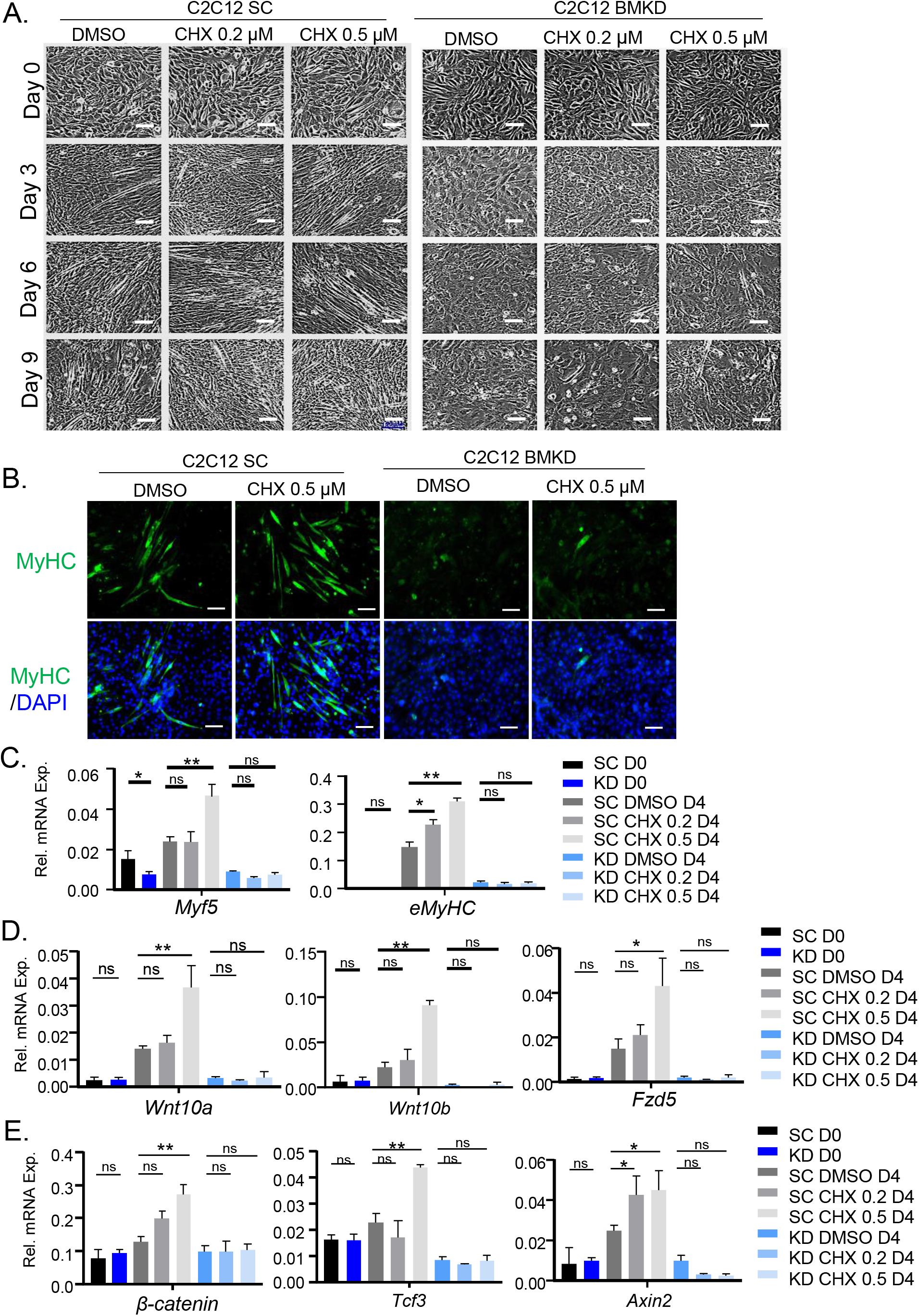
Clock-dependent effect of chlorhexidine on myogenesis. (A) Representative phase-contrast images of myogenic differentiation morphology at Day 0, 3, 6 and 9 in response to CHX treatment in C2C12 myoblasts with stable scrambled control shRNA (SC) or Bmal1 shRNA (BMKD). Scale bar: 100 μm. (B) Representative images of myosin heavy chain (MyHC) immunofluorescence staining at day 5 of myogenic differentiation in SC and BMKD myoblasts. Scale bar: 100 µm. (C, D) RT-qPCR analysis of myogenic gene expression (C), and genes of Wnt pathway (D) in SC and Bmal1 BMKD C2C12 myoblasts in response to CHX treatment before and at day 4 of C2C12 myogenic differentiation. Data are presented as Mean ± SD of n=3 replicates with two independent repeat experiments. *,**: p<0.05 or 0.01 CHX vs. DMSO by Student’s t test.

### Chlorhexidine promotes myoblast proliferation and migration

Muscle myogenic progenitors provides the major cellular source for regeneration, and its activation, proliferative expansion and migratory activity in response to injury are key processes involved in regenerative repair (53-55). We thus determined potential effects of CHX on the proliferative and migrative activities of myoblasts. As indicated by EdU incorporation, CHX at both concentrations tested induced ∼40% higher proliferation (**Fig. 7A** and **7B**) as compared to DMSO control. Furthermore, we included another clock modulator, the Rev-erbα antagonist SR8278, as a positive control based on our previous study of its effect on promoting myoblast proliferation (33). Comparison of the effect of CHX with SR8278 revealed that CHX at 0.2 µM is more efficacious to promote myoblast proliferation than SR8278 at 2.5 µM (**Fig. 7B**). A similar degree of augmented proliferation by CHX was observed in U2OS cells (**Fig. S4A & S4B**). Furthermore, the dose-dependent effect of CHX on promoting U2OS proliferation is dependent on a functional clock regulation, as this activity is abolished in cells with *CLOCK* gene silencing (**Fig. S4C & S4D**). Using a wound healing assay, treatment of CHX ranging from 0.2 to 1 µM significantly promoted area closure in C2C12 myoblasts when examined at 8 and 24 hours as compared with DMSO control, indicative of enhanced rate of cellular migration (**Fig. 7C** & **7D**). Together these analyses indicate that in addition to its effect on promoting myogenic differentiation, CHX also augmented the proliferative and migratory activities of myoblasts that may contribute to regenerative repair.

**Figure 7.**
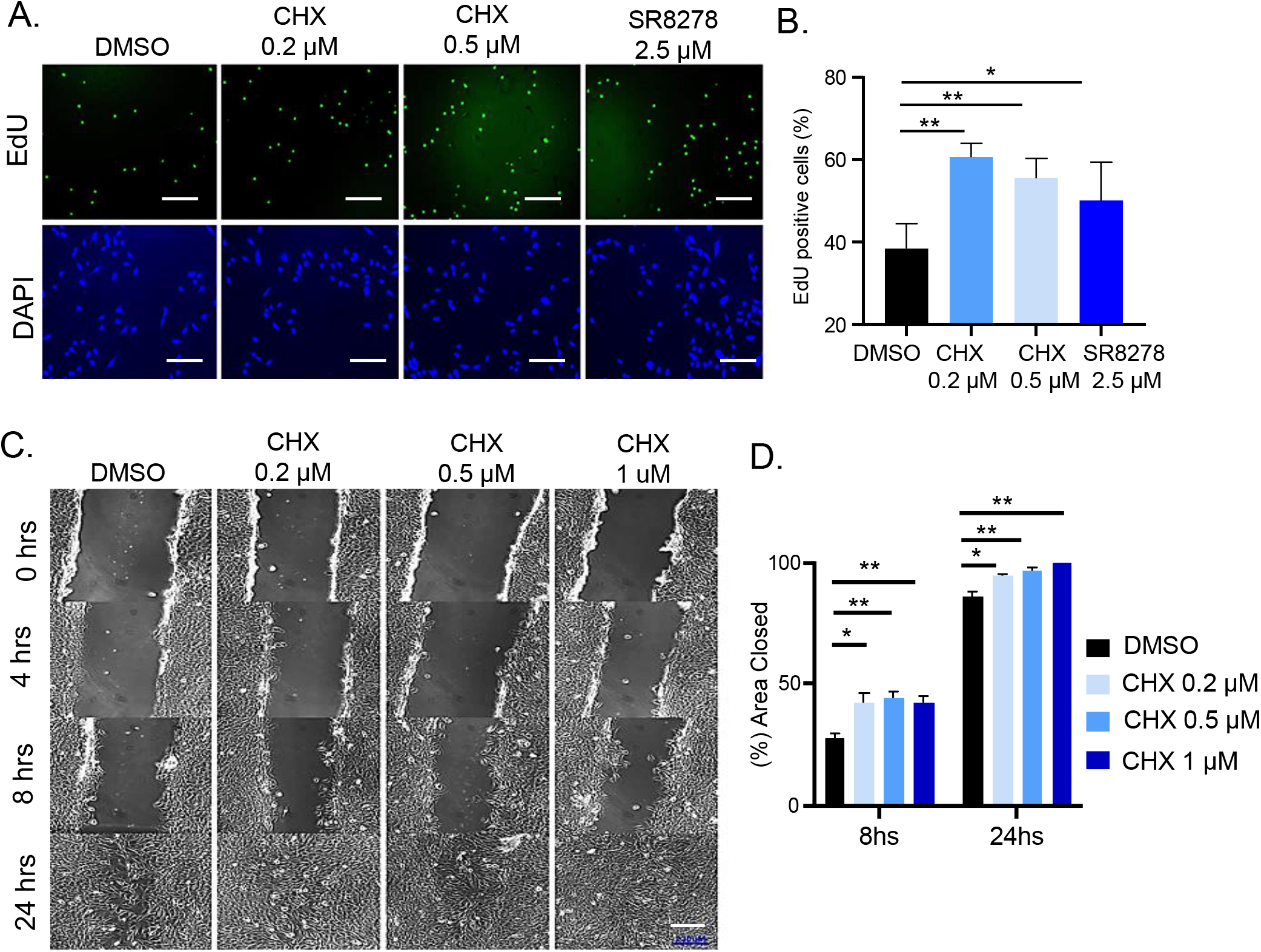
Effect of chlorhexidine on promoting myoblast proliferation and migration. (A, B) Representative images of EdU staining of C2C12 myoblasts with CHX or SR2878 treatment for 24 hours (A), with quantitative analysis (B). 10 representative fields were used for quantitative analysis. *,**: p<0.05 and 0.01 treated vs. DMSO by Student’s t test. Scale bar: 100 µm. (C, D) Representative images of wound healing assay of C2C12 myoblasts for 24 hours with CHX treatment at indicated concentrations (C) with quantitative analysis (D). 10 representative fields for each concentration. *,**: p<0.05, 0.01 and 0.001, CHX vs. DMSO by Student’s t test. Scale bar: 230 µm.

## Discussion

Circadian clock disruption leads to the development of obesity, insulin resistance and various types of cancer (56, 57). Thus, re-enforcing clock regulation may yield metabolic benefits and have potential for anti-cancer therapies. Despite current interests in developing clock modulators for disease applications, compounds directly modulate CLOCK activity has yet to be identified. Our screen specifically targeting the CLOCK protein led to the identification of CHX as a novel circadian clock-activating molecule that augments myoblast differentiation, proliferation and migratory activity. Given the role of coordinated circadian clock control in muscle stem cell behavior required for regenerative repair and protection against dystrophic muscle damage, the pro-myogenic properties of CHX may have therapeutic applications in dystrophic diseases or muscle-wasting conditions.

We leveraged a powerful high throughput in-silico screening platform based on molecular docking analysis of ∼300K compounds of NCI-DTP and FDA libraries to identify molecules with a structural fit for a hydrophobic pocket within the PAS-A domain of the CLOCK protein. Secondary functional screening for circadian clock properties of the top hits through virtual screen identified compounds with clock-modulatory activities, with further biochemical validation for effects on CLOCK/Bmal1-mediated gene transcription and heterodimer interaction. CLOCK forms an obligatory dimer with Bmal1 to activate transcription, and the PAS-A/B domains are structural elements mediating CLOCK and Bmal1 interaction. The docking of CHX with the CLOCK crystal structure revealed interactions with several key residues that forms a deep hydrophobic pocket within the PAS-A domain. As CHX displayed clock-activating effect in Per2-Luc reporter cell line, we postulated that its binding within the CLOCK hydrophobic pocket may enhance interaction affinity with Bmal1 to promote CLOCK/Bmal1-mediated transcription activation. Co-immunoprecipitation indeed revealed increased CLOCK and Bmal1 association in cells treated with CHX, with induction of clock genes consistent with its clock-activating activity. As a lead candidate identified from our screening, CHX chemical structure could be utilized as a basis for medicinal chemistry optimization to obtain derivatives with improved efficacy toward clock-modulation.

Recent endeavors to identify clock-modulating compounds are focused on Rev-erbα, RORα and Cry. Rev-erbα/β and RORs are ligand-binding nuclear receptors, and are therefore readily “druggable” by targeting the well-characterized ligand-binding pockets (23, 24, 38, 58, 59). Cry modulators identified so far function by inhibiting its proteosome-mediated degradation that stabilizes the protein and its clock-repressor function(14, 25, 60). To date, clock-modulating compounds that directly target CLOCK and Bmal1, the key transcription activators that drive core clock transcription feed-back loop, have not been discovered. This current study is the first report of a CLOCK protein-targeting molecule with biological activity in promoting myogenic differentiation. We also provide experimental evidence for the targeting specificity of CHX, with the absence of CHX activity in clock mutant and loss of its pro-myogenic effect in clock-deficient myoblasts. Mutating Cysteine 267 of CLOCK protein abrogated CHX-induced increased CLOCK/Bmal1 association, demonstrating the targeting specificity of the compound, while direct binding affinity of CHX with CLOCK remains to be determined in future investigations. Additional studies are needed to test the specificity of CHX, particularly against proteins with structural homology with CLOCK. Nonetheless, by testing CHX activity using CLOCK mutant, Bmal1-deficient myoblasts and CLOCK knockdown U2OS cells, our findings indicate that clock gene induction, myogenesis-enhancing activities and proliferative effects of CHX are dependent on a functional clock.

CHX is a cationic bisbiguanide molecule capable of binding to bacterial cell wall with bacteriostatic or bactericidal efficacy against a broad range of bacteria (61). It is commonly used as a topical skin antiseptic agent and an ingredient of mouth wash due to its anti-plaque and anti-gingivitis activity (62, 63), while known to bind to proteins on skin or mucous membranes with limited systemic absorption. Potentially due to its chemical properties, we found that, at concentrations higher than 1-2 µM, CHX may exhibit toxicity to cells, although distinct cell types appear to have differing sensitivity to CHX and is dependent on treatment duration. Dose response analysis of bioluminescence activity using Per2-Luc U2OS cells revealed a relatively broader range (0.2-2 µM) than functional assays for its pro-myogenic activities in C2C12 myoblasts (0.2-0.5 µM). We confined most of our experiments within a lower concentration range to avoid potential adverse effects, as compared to similar studies for clock modulator compound screening that typically uses 0.5-10 uM (50) or investigation of biological functions (24). Nonetheless, there could be potential clock-independent off-target effects of CHX, particularly its pro-myogenic activities in myoblasts. Despite its significant pro-myogenic action, the observed clock-modulatory effects of CHX on period lengthening was relatively moderate, which could be due to its mechanism of action by promoting CLOCK interaction with Bmal1. As our findings indicate that CHX augments the affinity of these interacting partners, there could be a limitation posed by the binding kinetics of these proteins in vivo, and thus, not as efficacious as potential clock inhibitors that disrupt the endogenous interaction. In line with this notion, clock-inhibitory molecules we discovered from this screen indeed are more potent in inhibiting Bmal1/CLOCK interaction and altering period lengthening (unpublished data). These observations collectively highlight the need for optimization of CHX chemical structure in order to develop new efficacious clock-activating molecules with pro-myogenic activities, while minimizing toxicity and off-target effects. Optimization of this lead compound may lead to development of novel agents for dystrophic muscle disease or metabolic disease applications.

Current pharmacological modulators of circadian clock components are mostly focused on metabolic and anti-cancer therapies (14, 19). The pro-myogenic properties of CHX was discovered based on our prior studies of circadian clock regulation in myogenesis. Both positive and negative arms of clock transcriptional loop modulate myogenic differentiation, with Bmal1 promoting whereas Rev-erbα inhibiting this process (6, 15, 16, 32, 33, 64). Notably, these regulations impact chronic dystrophic muscle damage in a preclinical animal model for Duchene Muscular Dystrophy, the dystrophin-deficient mdx mice (15, 16). Bmal1 function in promoting regenerative myogenesis is required to prevent dystrophic muscle damage (15), whereas loss of Rev-erbα ameliorates dystrophic pathophysiology (16). Moreover, pharmacological inhibition of Rev-erbα by an antagonist, SR8278, attenuated pathophysiological changes in the mdx model(65), providing the proof-of-principle validation for targeting clock components for dystrophic disease applications. In line with these prior findings, activation of clock by CHX in myoblasts resulted in enhanced differentiation and this effect is dependent on a functional clock. In addition, CHX augmented the proliferation and migration of myoblasts, implicating its potential utility to facilitate distinct stages of myogenic regenerative repair to ameliorate dystrophic disease. This study, together with our prior findings, suggest that clock modulators have potential utilities in muscle diseases, such as muscular dystrophy or muscle-wasting condition associated with chronic inflammation or aging. Future studies for in vivo testing of the efficacy and potential adverse effects in pre-clinical models of muscle diseases is needed.

Accumulating epidemiological and experimental studies collectively established that disruption of clock regulation leads to the development of obesity and insulin resistance (66). We previously reported that loss of Bmal1 resulted in obesity (37), and chronic shiftwork induced adipose tissue expansion with inflammation (67). The activity of CHX in promoting clock function, conceivably, could be applied to metabolic disease treatment. A number of small molecule clock modulators discovered to date have demonstrated efficacy in metabolic disease applications. Rev-erbα agonists, SR9009 and 9011, displayed anti-obesity and anti-hyperlipidemia actions (24, 58). Nobiletin, a ROR agonist, promotes mitochondrial oxidative capacity in skeletal muscle to protect against development of insulin resistance induced by a high fat diet challenge (39, 40). Therefore, it is conceivable that small molecules directly targeting CLOCK, such as CHX or its derivatives, may have potential utilities in obesity or diabetes. The metabolic activities of CHX awaits further studies. On the other hand, by activating clock, CHX displayed pro-myogenic effects while enhancing myoblast proliferation. CHX may have potential undesirable effects on cell proliferation that increases the risk for cancer, and we found CHX induced proliferation of U2OS cells. Developing new compounds that dampen the proliferative activity may reduce this adverse effect on cells that is a pre-requisite for future drug development efforts.

## Conclusions

In summary, our study is the first report to demonstrate the feasibility of a screening pipeline for discovering circadian clock modulators by targeting the CLOCK protein that has myogenic-regulatory properties. The discovery of CHX as a pro-myogenic molecule targeting the clock modulation may have potential applications in muscular dystrophy or related muscle remodeling processes involving muscle stem cells.

## Non-standard Abbreviations

CLOCK: Circadian Locomotor Output Cycles Kaput
Bmal1: Brain and Muscle Arnt-like Protein 1
PAS: Per-Arnt-Sim
ROR: RAR-related Orphan Receptor
CHX: chlorhexidine
MyHC: myosin heavy chain

## Declarations

### Ethics approval and consent to participate

Not applicable

### Consent for publication

Not applicable

### Availability of data and materials

All data generated and analyzed during this study are included in this published article and associated Supplementary Information files.

### Authors’ contributions

TK, WL and XX: data curation and investigation, formal analysis, manuscript editing; HL and DH: data curation and investigation; KM: conceptualization, formal analysis, project administration, manuscript writing and editing, and funding acquisition.

## Acknowledgements

We thank Drs. Steve Kay and Meng Qu at the University of Southern California for sharing the luciferase reporter cell lines used in this study, and Drs. Seung-Hee Yoo and Zheng Chen at University of Texas at Houston Health Science Center for providing the Per2-luciferase plasmid.

