## Supplemental Figures for "Targeted Screening and Identification of Chlorhexidine as a Pro-myogenic Circadian Clock Activator"

**Supplemental Figure S1.**

A.

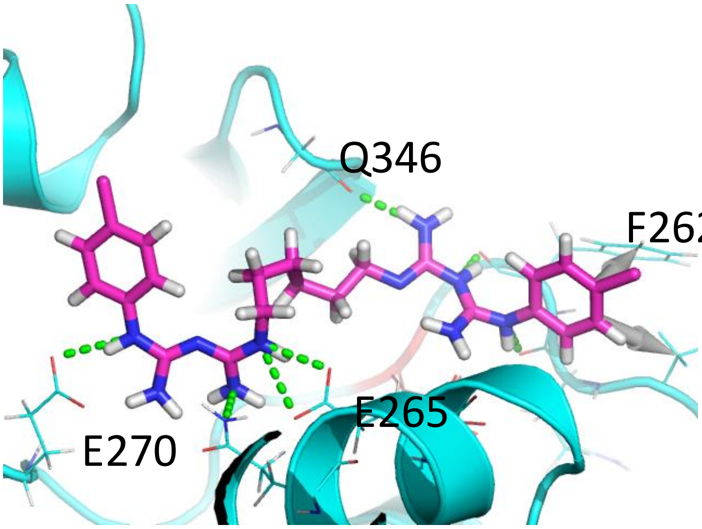


B.

| Predicted Distance/Interaction | CLOCK AA Residue Position |
|---|---|
| ≤3A | Q112 E116 A117 D119 Q261 F262 K264 E270 P295 G345 Q346 Q347 |
| ≤4A | P260 M266 |
| ≤6A | L113 L115 L118 L141 R256 I263 T268 E271 E274 P294 I296 G298 T343 K344 |
| Hydrogen Bond | E116 Q261 F262 E270 G345 Q112 |
| Hydrophobic interaction | PHE262, PRO260 |

12 Residues within 3A: Q112 E116 A117 D119 Q261 F262 K264 E270 P295 G345 Q346 Q347
2 Residues within 4A : P260 M266
14 Residues within 6A: L113 L115 L118 L141 R256 I263 T268 E271 E274 P294 I296 G298 T343 K344
6 unique residue for Hydrogen Bonds: E116 Q261 F262 E270 G345 Q112
2 Hydrophobic interaction pairs at 5A: PHE262, PRO260

**Figure S1. Chlorhexidine interactions with residues within the CLOCK protein hydrophobic pocket**. (A) Docking pose of CHX, represented as the pick stick molecule, on CLOCK hydrophobic pocket structure with interacting amino acid residues listed. (B) List of predicted CHX interactions with amino acid residues within the CLOCK hydrophobic pocket by all-around docking modeling.

**Supplemental Figure S2.**

A.

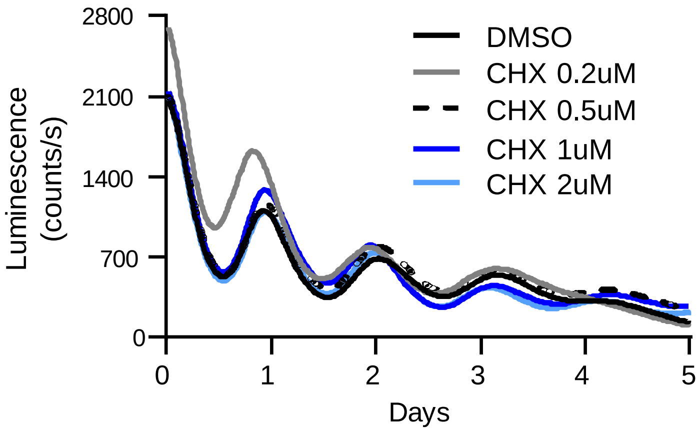


B.

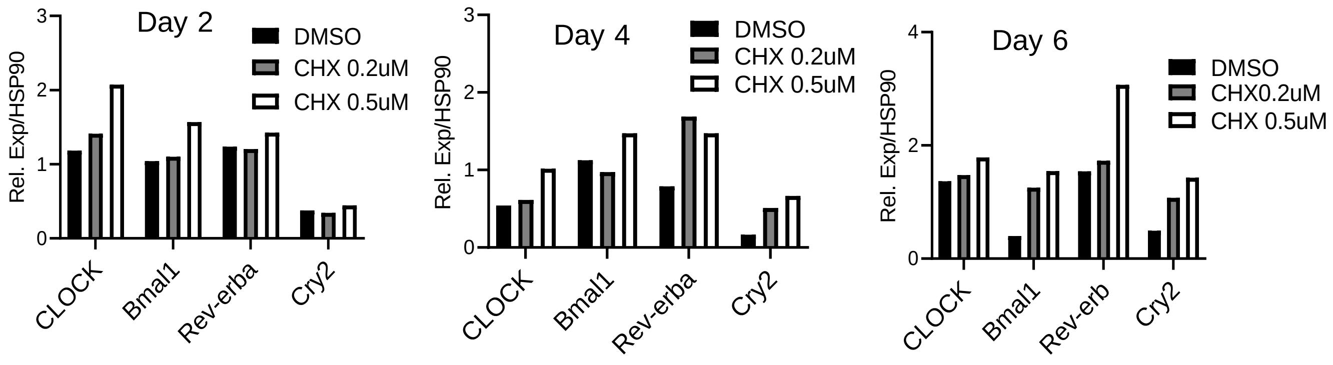


Figure. S2. (A) Original plot of average tracing of bioluminescence activity monitoring of Per2::dLuc U2OS reporter cell line for 6 days using Lumicycle, with Chlorhexidine treatment (CHX) at indicated concentrations as shown in baseline-adjusted plots. N=4/treatment. (B) Quantification of immunoblot analysis shown in Fig. 4A of Chlorhexidine effect on clock protein expression levels at day 2, 4 and 6 of myogenic differentiation.

**Supplemental Figure S3.**

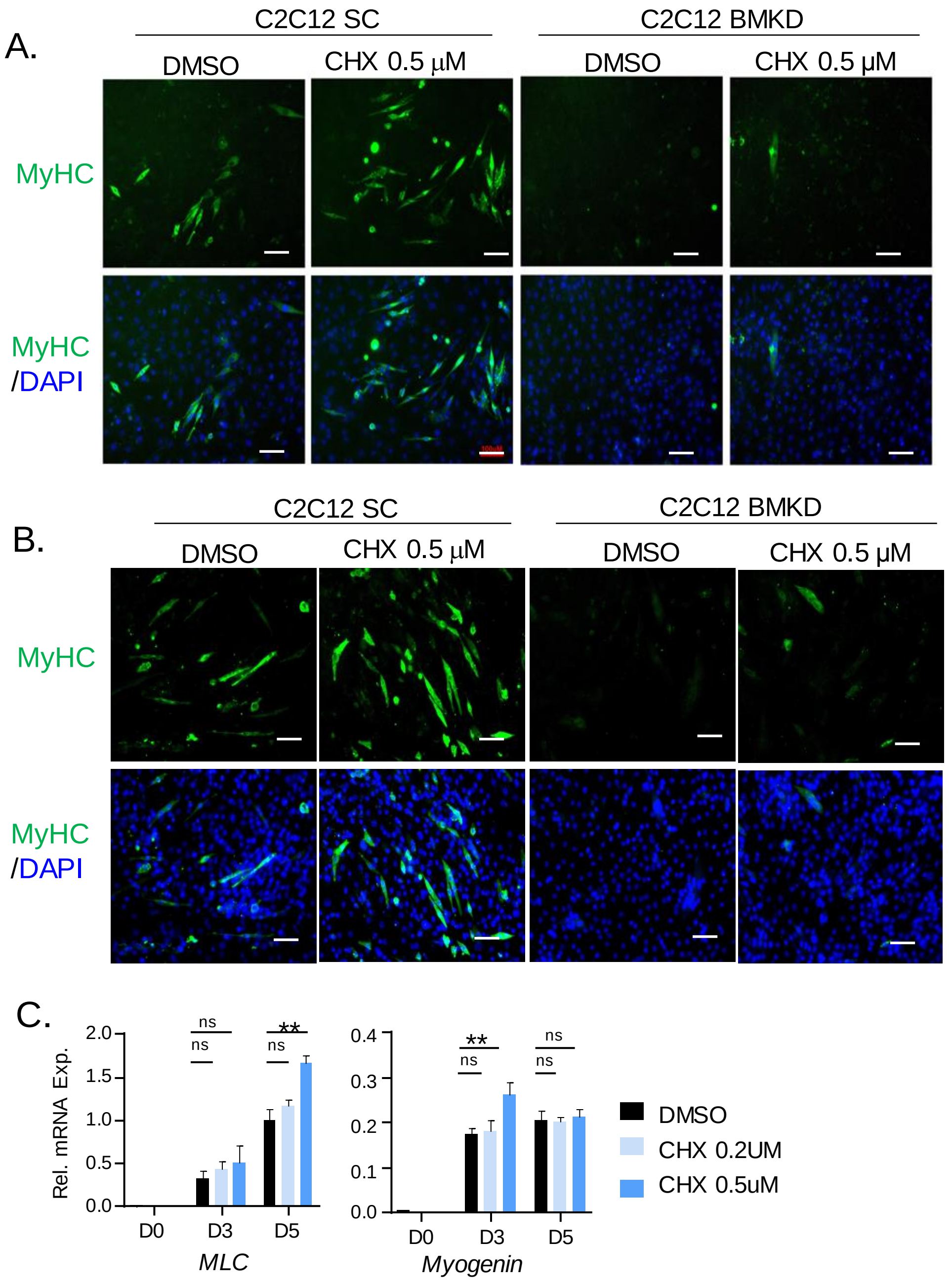


**Figure S3. Effect of Chlorhexidine on C2C12 myoblast differentiation.** (A, B) Representative images of immunofluorescence staining of myosin heavy chain (MyHC) at day 3 (A) and day 5 (B) of SC control and BMKD C2C12 myoblast differentiation at indicated CHX concentration. Scale bar: 100 μm. (C) RT-qPCR analysis of myogenic gene expression after CHX treatment for 24 hours at day 0, 3 and 5 of C2C12 myogenic differentiation. *MLC*: Myosin light chain. *,**: $p<0.05$ and 0.01 CHX vs. DMSO by Student's t test. N=3/group.

### Supplemental Figure S4.

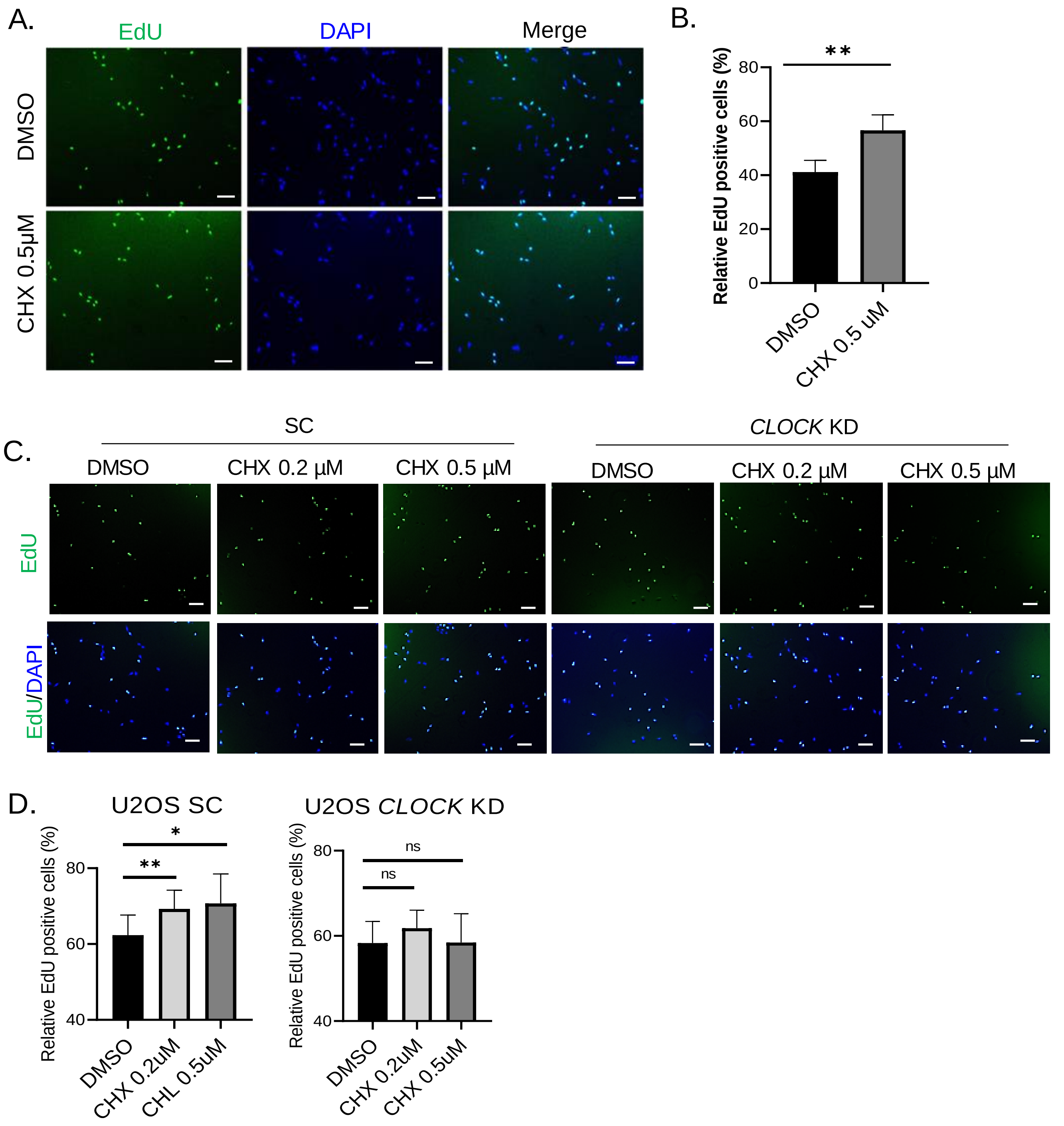


**Figure S4: Effect of Chlorhexidien on proliferation of U2OS cells.** (A, B) Representative images of EdU and DAPI staining of U2OS cells with DMSO or CHX treatment for 24 hours (A), with quantification (B). 3 repeats of 10 representative fields was used for quantitative analysis.. (C, D) Representative images of EdU and DAPI staining of U2OS with stable expression of scrambled control shRNA (SC) or *CLOCK* shRNA (*CLOCK* KD) treated with DMSO or CHX treatment for 24 hours (A), with quantification (B). 3 repeats of 10 representative fields was used for quantitative analysis. *,**: p<0.05 and 0.01, CHX vs. DMSO by Student's t test. Scale bar: 100 μm.

**Supplemental Figure S5:** Uncropped gels as for Figure 2E & 2F

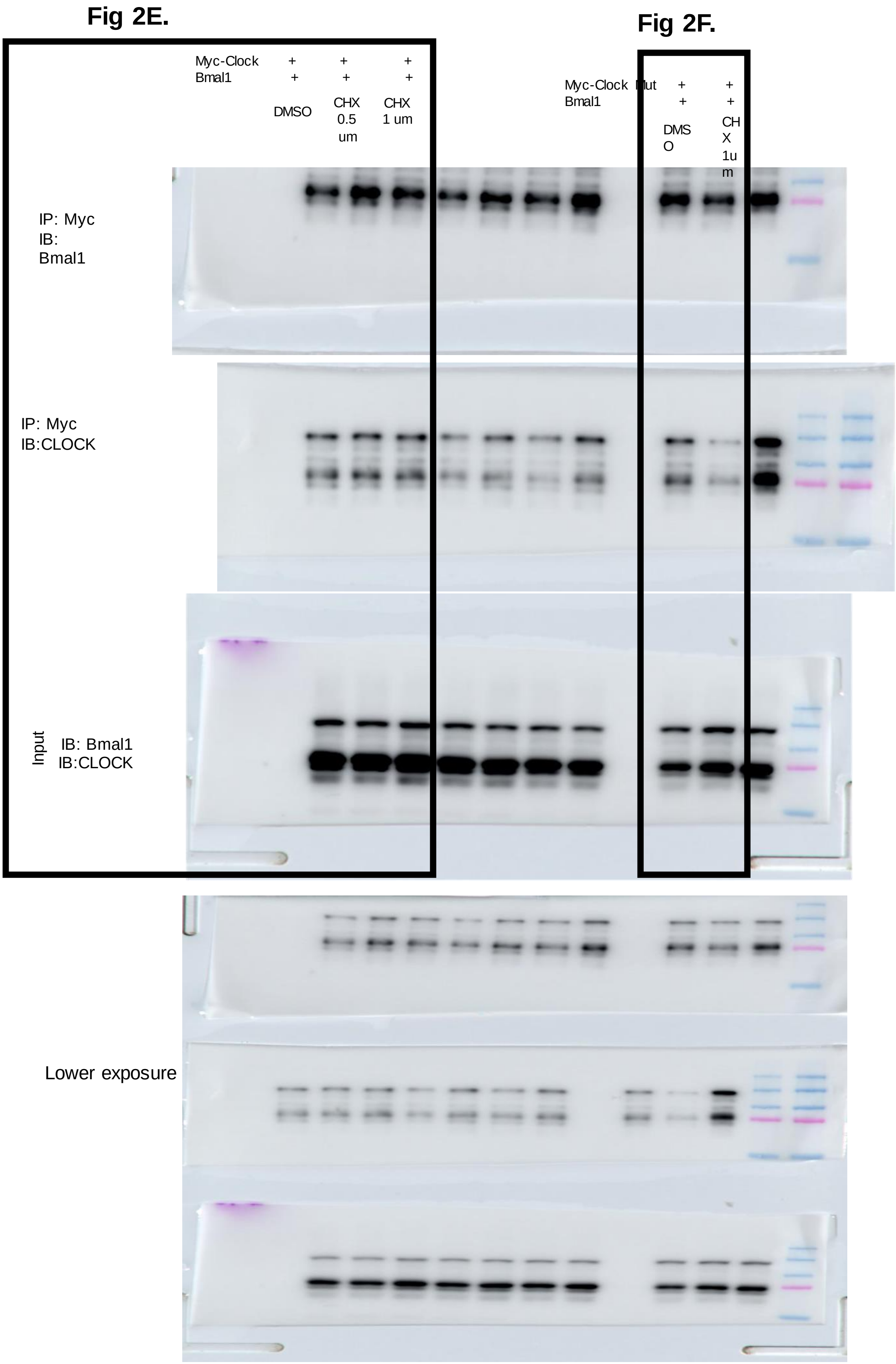

**Supplemental Figure S6:** Uncropped gels as for Figure 3A.

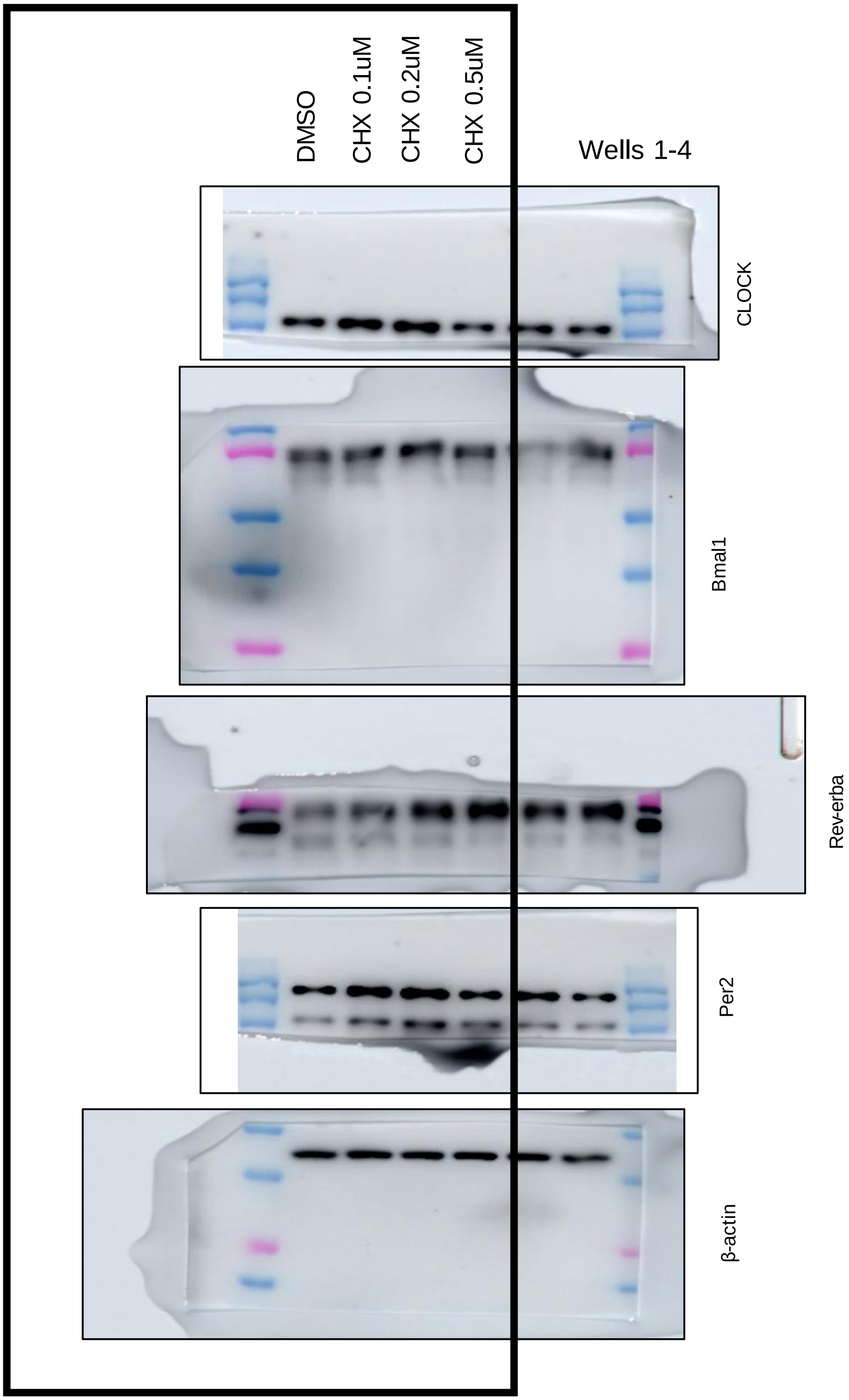

**Supplemental Figure S7:** Uncropped gels for Figure 4A

.

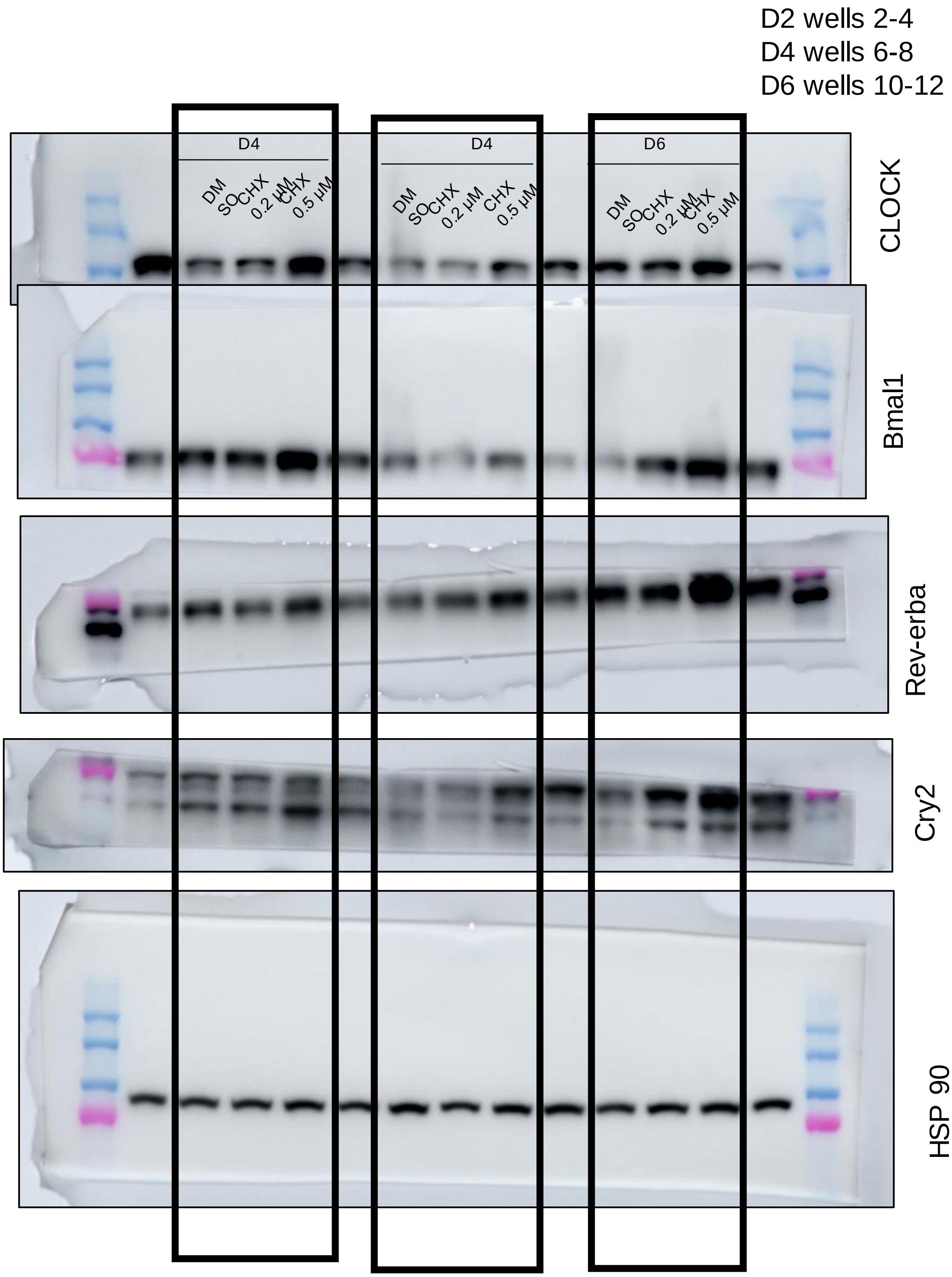

**Supplemental Figure S8:** Uncropped gels for Figure 5C.

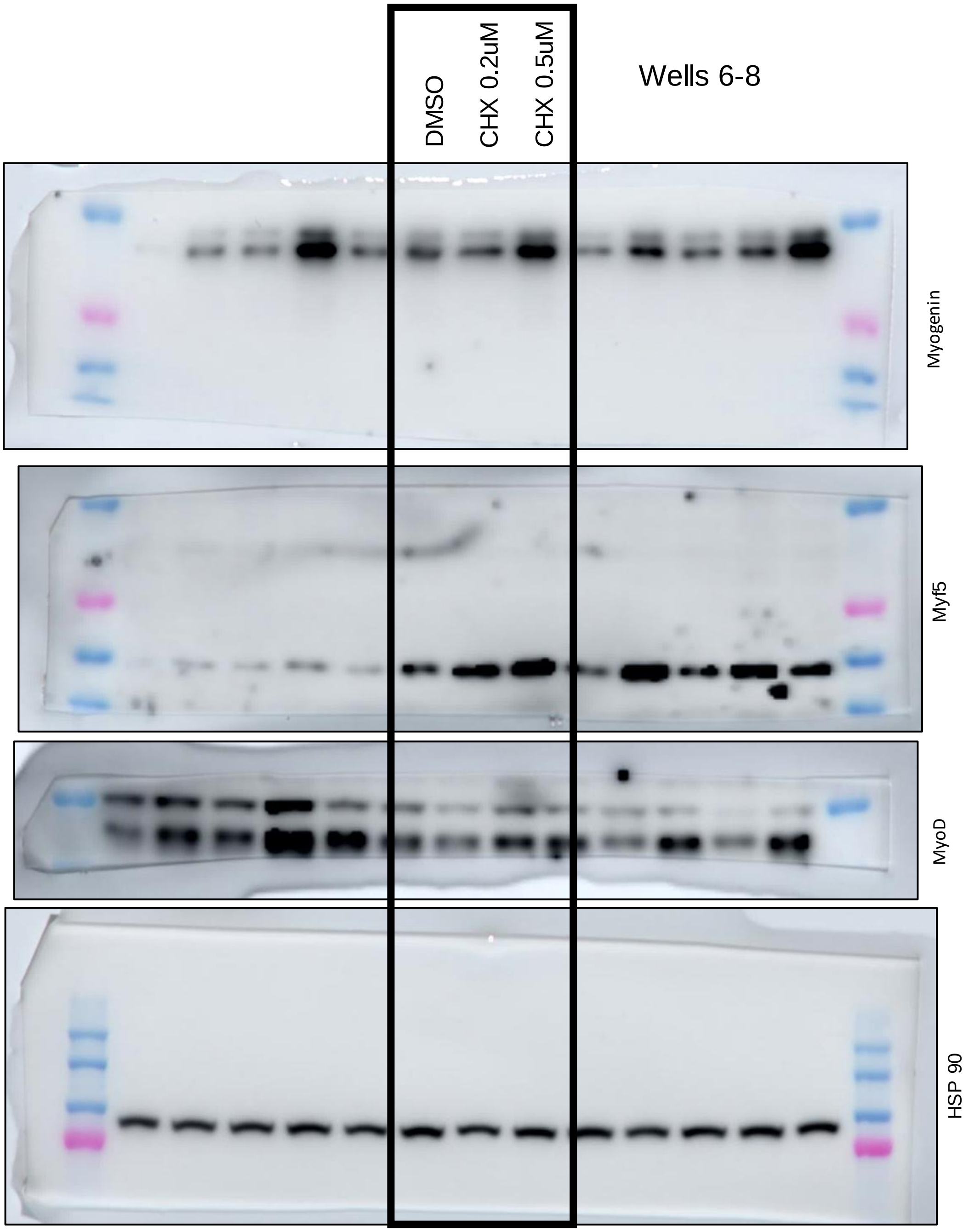

**Suppl. Table 1.** List of cell lines.

| Cell line | Tissue | Cell type | Mediu m | Source | Obs. |
|---|---|---|---|---|---|
| **U2OS** | Bone | Epithelial | DMEM | Dr. Steve Kay at University of Southern California | Per2-Luciferase reporter<br>Liu et al. PLoS Genet. 2008 Feb 29;4(2):e1000023. |
| **293A** | | Epithelial | DMEM | ATCC® CRL-1573 | |
| **C2C12** | Muscle | Myoblast | DMEM | ATCC® CRL-1772 | |
| **C2C12**<br>**SC control**<br>**Bmal1 KD** | Muscle | Myoblast | DMEM | Dr. Ke Ma at City of Hope | Bmal1 small hairpin RNA (shRNA) construct in pGIPZ vector (VGM5520–99941526, VGM5520–99342254, VGM5520–99211363) from Open Biosystems<br>Chatterjee et al. J Cell Sci 2013 May 15;126(Pt 10):2213-24. |

**Suppl. Table 2.** List of primer sequences for RT-qPCR analysis.

| Primer | Organism | Sequence |
|---|---|---|
| **36B4-F** | Human | AACATGCTCAACATCTCCCC |
| **36B4-R** | Human | CCGACTCCTCCGACTCTTC |
| **Bmal1-F** | Human | CGGAGTCGATGGTTCAGTTT |
| **Bmal1-R** | Human | CTTCCAGGACGTTGGCTAAA |
| **Clock-F** | Human | TGCGAGGAACAATAGACCCAA |
| **Clock-R** | Human | ATGGCCTATGTGTGCGTTGTA |
| **DBP-F** | Human | CTGATCTTGCCCTATCAAGCATT |
| **DBP-R** | Human | CGATGTCTTCGAGGGTCAAAG |
| **Per1-F** | Human | CCAGGATGTGGGAGTCTTCTAT |
| **Per1-R** | Human | ACGGACTTCTCCTGGGTAAA |
| **Per2-F** | Human | TGCAGTGGAGCAGATTCTTT |
| **Per2-R** | Human | GGTGGTAGCGGATTTCATTCT |
| **Cry1-F** | Human | CCTGTGGATTAGTTGGGAAGAA |
| **Cry1-R** | Human | CTACAAGACAGCCACATCCAA |
| **Cry2-F** | Human | GGGTCCGGGTATTTGATGAG |
| **Cry2-R** | Human | GTGGAAGAACTGCTGGAAGA |
| **NR1D1-F** | Human | TGGACTCCAACAACAACACAG |
| **NR1D1-R** | Human | GATGGTGGGAAGTAGGTGGG |
| **NR1D2-F** | Human | TTTAGTGGCATGGTTCTACTGTG |
| **NR1D2-R** | Human | AGCCTTCGCAAGCATGAACT |
| **36B4-F** | Mouse | CGCTTTCTGGAGGGTGTCCGC |
| **36B4-R** | Mouse | TGCCAGGACGCGCTTGTACC |
| **Bmal1-F** | Mouse | ACTCACCGTGCTAAGGATGGCTGT |
| **Bmal1-R** | Mouse | CTCGGAGACAAAGAGGATCTTCCCTCG |
| **Clock-F** | Mouse | TTGCTCCACGGGAATCCTT |
| **Clock-R** | Mouse | GGAGGGAAAGTGCTCTGTTGTAG |
| **DBP-F** | Mouse | CCACCGCGCAGGCTTGACAT |
| **DBP-R** | Mouse | ACAGGGCGAGATCAGCGGGA |
| **Per1-F** | Mouse | CTGCCATGGAGGAAGAAGAG |
| **Per1-R** | Mouse | AGCTGGGGCAGTTTCCTATT |
| **Per2-F** | Mouse | ATGCTCGCCATCCACAAGA |
| **Per2-R** | Mouse | GCGGAATCGAATGGGAGAAT |
| **Cry1-F** | Mouse | CTGGCGTGGAAGTCATCGT |
| **Cry1-R** | Mouse | CTGTCCGCCATTGAGTTCTATG |
| **Cry2-F** | Mouse | TGTCCCTTCCTGTGTGGAAGA |
| **Cry2-R** | Mouse | GCTCCCAGCTTGGCTTGA |
| **NR1D1-F** | Mouse | TGGCATCCGGTGCACTGCAG |
| **NR1D1-R** | Mouse | CCCTCCAGAAGGGTAGCACGCT |
| **NR1D2-F** | Mouse | GGAGTTCATGCTTGTGAAGGCTGT |
| **NR1D2-R** | Mouse | CAGACACTTCTTAAAGCGGCACTG |
| **eMyHC-F** | Mouse | AAAAGGCCATCACTGACGC |
| **eMyHC-R** | Mouse | CAGCTCTCTGATCCGTGTCTC |
| **MLC-F** | Mouse | GCAACAGGAGGACTTCAAGGAGGC |
| **MLC-R** | Mouse | ATTGGTGCCCAGAGCCCGGA |
| **Myf5-F** | Mouse | AGCTGCTGAGGGAACAGGTGGA |
| **Myf5-R** | Mouse | ATTCAGGCATGCCGTCAGAGCA |
| **MyoD-F** | Mouse | AACCCAGACCCCGTCTCCCG |
| **MyoD-R** | Mouse | GGGCGTGAAGAACCAGGGGC |
| **Myogenin-F** | Mouse | GTCCCAACCCAGGAGATCATT |
| **Myogenin-R** | Mouse | AGTTGGGCATGGTTTCGTCT |

**Suppl. Table 3.** List of primary and secondary antibodies for immunoblot analysis.

| Protein | Manufacturer | Catalog # | Dilution (fold) |
|---|---|---|---|
| **βActin** | Invitrogen | MA5-15739 | 1:10000 |
| **HSP90** | Cell Signaling Technology | 4847S | 1:5000 |
| **Bmal1** | Cell Signaling Technology | 14020S | 1:5000 |
| **CLOCK** | Cell Signaling Technology | 5157S | 1:4000 |
| **NR1D1** | Proteintech | 14506-1-AP | 1:3000 |
| **Cry2** | Proteintech | 13997-1-AP | 1:4000 |
| **Per2** | Proteintech | 67513-1-Ig | 1:4000 |
| **MyoD** | Santa Cruz Biotechnology | sc377460 | 1:2000 |
| **Myogenin** | Santa Cruz Biotechnology | sc52903 | 1:2000 |
| **Myf5** | Santa Cruz Biotechnology | Sc518039 | 1:2000 |
| **Myc** | Cell Signaling Technology | 2278 | 1:5000 |
| **PAX7** | DSHB | - | 1:500 |
| **Goat anti Rabbit** | Bio Rad | 5213-2504 | 1:5000 |
| **Goat anti Mouse** | Bio Rad | STAR 207 | 1:5000 |

**Suppl. Table 4.** List of primary and secondary antibodies for immunofluorescence staining.

| Protein | Manufacturer | Catalog # | Dilution (fold) |
|---|---|---|---|
| **MyHC** | Developmental Studies Hybridoma Bank | MF 20 | 1:50 |
| **Alexa Fluor 488 goat anti-mouse IgG** | Thermo Fisher Scientific | A-11001 | 1:1000 |
| **Alexa Fluor 594 goat anti-mouse IgG** | Thermo Fisher Scientific | A-11005 | 1:1000 |
